# Integrative prediction of gene expression with chromatin accessibility and conformation data

**DOI:** 10.1101/704478

**Authors:** Florian Schmidt, Fabian Kern, Marcel H. Schulz

## Abstract

**Background:** Enhancers play a fundamental role in orchestrating cell state and development. Although several methods have been developed to identify enhancers, linking them to their target genes is still an open problem. Several theories have been proposed on the functional mechanisms of enhancers, which triggered the development of various methods to infer promoter enhancer interactions (PEIs). The advancement of high-throughput techniques describing the three-dimensional organisation of the chromatin, paved the way to pinpoint long-range PEIs. Here we investigated whether including PEIs in computational models for the prediction of gene expression improves performance and interpretability.

**Results:** We have extended our Tepic framework to include DNA contacts deduced from chromatin conformation capture experiments and compared various methods to determine PEIs using predictive modelling of gene expression from chromatin accessibility data and predicted transcription factor (TF) motif data. We found that including long-range PEIs deduced from both HiC and HiChIP data indeed improves model performance. We designed a novel machine learning approach that allows to prioritize TFs in distal loop and promoter regions with respect to their importance for gene expression regulation. Our analysis revealed a set of core TFs that are part of enhancer-promoter loops involving YY1 in different cell lines.

**Conclusion:** We show that the integration of chromatin conformation data improves gene expression prediction, underlining the importance of enhancer looping for gene expression regulation. Our general approach can be used to prioritize TFs that are involved in distal and promoter-proximal regulation using accessibility, conformation and expression data.

## Introduction

Understanding the processes involved in gene regulation is an important endeavour in computational biology. Key players in gene regulation are transcription factors (TFs), DNA binding proteins that are essential in regulating transcriptional processes. They are important in establishing and maintaining cellular identity and their dysfunction is related to several diseases [1].

TFs bind to promoters of genes, which are in close proximity to their transcription start site (TSS) and to enhancers, regulatory regions that can be several thousand base pairs away from the regulated gene [2]. Since enhancers have been described for the first time in 1981 by *Banerji et al.* [3], numerous studies shed light on their functional role.

For example, enhancers were shown to be essential in cell differentiation [4]. Also, it has been reported that mutations occurring in enhancer regions, can not only lead to changes in gene expression [5, 6] but can also increase the probability to contract certain diseases, for instance *Hirschsprung’s disease*[7]. These effects are likely to be caused by an altered binding of TFs due to SNPs occurring in enhancer sequences [2, 8, 9]. To understand the function of enhancers, a crucial step after identification of putative enhancer regions is to link them to their target genes.

Recently, considerable progress has been made in identifying putative enhancer regions: In the past decade, many epigenetic datasets have been generated in consortia like ENCODE [10], Blueprint [11], and Roadmap [12]. Histone Modifications, especially *H3K27ac* and *H3K4me1*, have been used in unsupervised computational approaches, such as ChrommHMM [13], EpiCSeg [14], or REPTILE [15] to highlight putative enhancers regions genome-wide.

Also (semi-)supervised methods, e.g. McEnhancer [16], EnhancerDBN [17], or DECRES [18], relying on experimentally validated enhancer regions used as training data have been proposed. Furthermore, it was shown that DNase-hypersensitive sites (DHSs) are good candidate sites for TF-binding [19, 20] and that DNase1-seq signal is also predictive for gene expression [20, 21]. Thus DHS sites, which are not located nearby promoters can be considered as candidate enhancer regions. However, it is still a fundamental biological question how enhancers interact with their potentially distantly located target genes. The most prevalent hypothesis is that enhancers are brought to close proximity to their target genes by chromosomal re-organisation and DNA-looping. This hypothesis is known as the *looping* model. It is opposing the so-called *scanning* model, which states that an enhancer is usually regulating only its nearest active promoter [22]. Experimental evidence could be found for both models [2], hence it is likely that both mechanisms are occurring in nature.

Inspired by these models, several experimental and computational methods have been proposed to link enhancers to their target genes. Following the *scanning* model, two approaches are common in the field: [1] *window* based linkage and [2] *nearest gene* linkage. In the window based approach, a gene is associated to regulatory regions that are located within a defined genomic region around this gene [23, 24]. Alternatively, in the nearest gene approach, an enhancer is only associated to its nearest gene [25]. To reduce false positive assignments, the *nearest gene* linkage is also often coupled to a correlation test between epigenetic signals in the enhancer and the expression of the candidate gene [26].

While approaches like FOCS [27] or STITCHIT [28] offer the linkage of regulatory elements on a gene-specific level, these methods require the availability of large data sets for the considered species and tissues, which is generally not the case. In practice, the established *window* and *nearest gene* based linkage paradigms are still being used [25]. However, the drawback of those approaches is that they do not include long range enhancer-gene interactions, as proposed by the *looping* model. These have been experimentally determined using for example Fluorescence In Situ Hybridization (FISH), via the identification of enhancer RNAs (eRNAs) and their correlation to target genes, or via 3C-based high-throughput methods, for instance HiC, Capture-HiC and HiChIP [29]. Especially the development of such high-throughput methods to analyse the 3D organisation of the genome enables us to determine genome-wide DNA contacts [30]. Detailed analyses of individual genes, e.g. the *β-globin gene* showed that multiple contacts occur simultaneously at one genomic loci and also overlap with DHSs [31]. It was shown that loops are established by Cohesin, Mediator complexes, and CTCF, which is known to act as an insulator protein. By performing genome-wide chromatin conformation capture experiments, it is possible to segment the genome in multiple topological associating domains (TADs). It was shown before that there is more intra-TAD interaction among genes and enhancers than between TADs [32]. To mine this information, a wealth of tools have been published. A detailed overview is provided in Yao *et al* [2]. Despite the availability of such data, it has not yet been integrated into computational methods inferring gene expression using experimentally or computationally determined TF binding events. Because of the tissue specificity of enhancers, including PEIs might augment the interpretability of such models and thereby lead to novel biological insights.

Here, as a follow-up to our previous investigations [24, 33], we introduce an extension of the Tepic framework to account for PEIs inferred from chromatin conformation capture experiments. As a baseline, we illustrate that both *window* and *nearest gene* annotation approaches are not well suited to account for enhancer activity. Our new Tepic module extends a promoter-centric window by including far away genomic loci deduced from HiC and HiChIP data. While both HiC and HiChIP interactions improve the gene expression prediction models, we observe a greater improvement with HiChIP data, an effect for which we outline several reasons. Furthermore, we illustrate that a distinct consideration of TF binding events in promoter and potentially far away enhancer regions allows for a fine grained interpretation and analysis of transcriptional regulation through TFs.

## Materials and methods

### Data and Preprocessing

In this study we used gene expression quantified from RNA-seq data and DNase1-seq data for the cell lines K562, GM12878, IMR90, HUVEC, HCT116, Jurkat, and HeLa. All data is obtained from ENCODE, corresponding accession numbers are provided in Sup. Tab. 1. Except for Jurkat, where gene expression estimates were quantified with SALMON (version 0.8.2) using default parameters, gene expression estimates were directly downloaded from ENCODE. DHS sites have been identified using the peak caller JAMM [34] (version 1.0.7.2), with default parameters configured. All peaks passing the automated filtering of JAMM are considered. Sup. Tab. 2 lists the number of identified peaks per cell line.

Furthermore, we obtained HiC data for K562, GM12878, IMR90, HUVEC, and HeLa from Rao *et al* [30]. Specifically, we used the loop files as provided by the Lieberman-Aiden group, which were extracted from raw HiC contact matrices using the HiCCUPS algorithm [30]. In case of the HiC datasets used in this work the loops are of 5kb, 10kb, and 25kb resolution, respectively. A loop is defined as a pair of genomic loci that are in arbitrary genomic distance from each but, at the same time, are in close spatial proximity. In the following, the HiC resolution called *All* refers to loops of an arbitrary resolution, as this corresponds to a more conservative approach where we collect all available loops. For reasons of simplicity, inter-chromosomal loops, which resemble a less frequent type of contacts, are excluded. Sup. Tab. 3 provides an overview on the HiC data considered in this work.

Additionally, we use processed HiChIP data (Sup. Tab. 4) in which the TF YY1 was targeted in Jurkat, HCT116, and K562 cells generated by Weintraub *et al.* [35]. The data has a resolution of 5kb. All data was obtained for the hg19 reference genome using gene annotation version 19 from GENCODE [36].

We obtained chromatin state segmentations, containing 15 states generated with ChromHMM [13], for K562, GM12878, IMR90, HUVEC, and HeLa from ENCODE. As there was no ChromHMM annotation available for Jurkat, we approximate this using a Roadmap ChromHMM track for CD4+ CD25− Th Primary Cells (E043). We focus on the promoter states *TssA*(1) and *TssAFlnk* (2), as well as on the enhancer states *EnhG*(6), *Enh*(7), *BivFlnk* (11), and *EnhBiv* (12). ENCODE accession numbers are provided in Sup. Tab. 1.

### Establishing Promoter-Enhancer interactions (PEIs)

We apply three different strategies to come up with PEIs from DNase1-seq data: (1) A window based annotation, (2) a nearest gene-based linkage, and (3) a window based annotation that incorporates HiC or HiChIP data. An illustration of the PEI linkage methods is shown in Fig. 1.

**Figure 1:**
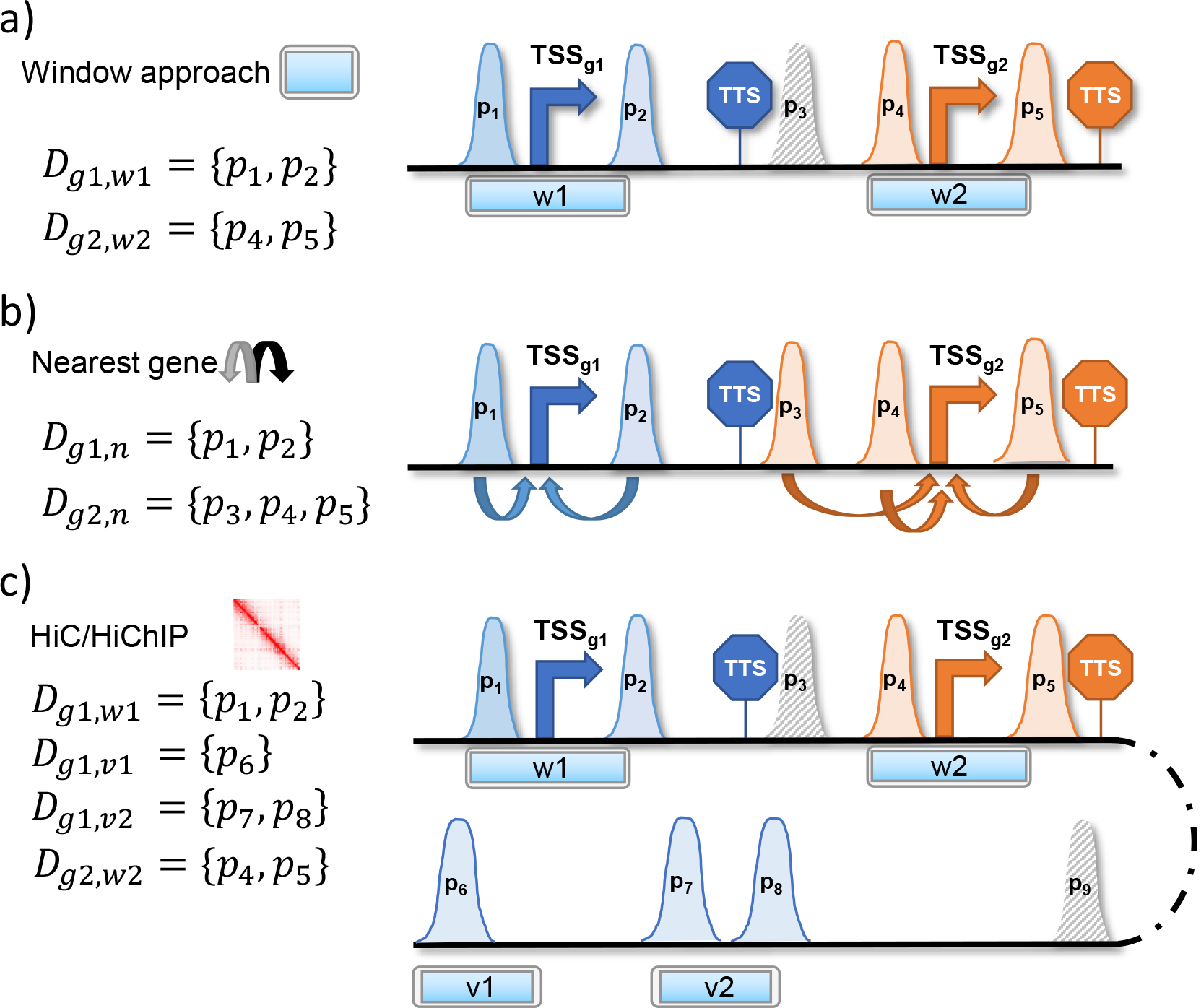
Assignment of DNase1-seq peaks to genes. The different setups are illustrated for two genes *g*1 and *g*2. The color code of peaks and the border color of segments indicate to which gene a peak is assigned. Peaks with a striped filling are not assigned to any gene. a) In a window based annotation, peaks are linked to a gene if they are located within a window *w* centered at the 5’ transcription start site (TSS) of a gene of interest. *D*_*g*1, *w*1_ denotes the set of all DHSs overlapping window *w*1 centered around the promoter of gene *g*1. b) Peaks are linked to the nearest gene, defining nearest as the gene with the closest TSS in linear genomic distance. Here, *D*_*g*1, *n*_ refers to the set of all DHSs linked to gene *g*1 following the nearest gene approach. c) Using HiC or HiChIP, secondary windows *v*_*i*_ covering the distal regions linked to the TSS are considered in addition to the TSS window. For gene *g*1, two additional windows, *v*1 and *v*2, are considered, yielding the additional peak sets *D*_*g*1, *v*1_, and *D*_*g*1, *v*2_.

We use the following notation throughout the article: Considering a DHS site 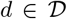, where 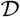 is the set of all DHS sites, we denote the length of *d* with *l*(*d*) and the DNase1-seq signal in *d* with *s*(*d*). We aggregate neighbouring genomic positions, which are assigned the same chromatin state from ChromHMM into one *segment m*, representing a distinct ChromHMM state. The set of all considered segments is denoted with 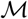.

Here, we compute three different features for each gene *g*: (1) total peak length *pl*_*g*_, (2) summarized peak count *pc*_*g*_, and (3) aggregated peak signal *ps*_*g*_ [33]:

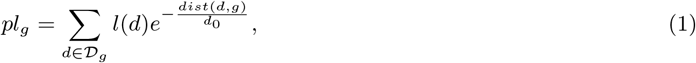

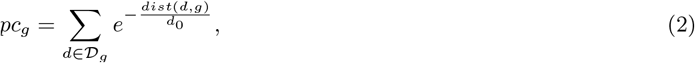

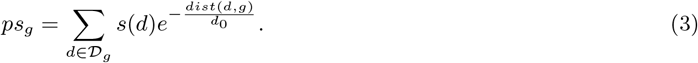

The genomic distance of *d*, *c*_*t*_, or *m* to a specific gene *g* is denoted with *dist*(*d*, *g*). The genomic distance is measured from the centre of the peak to the most 5′-TSS of *g*. Using an exponential decay formulation proposed by *Ouyang et al.* [23], each peak is weighted by the distance to its linked gene. The parameter *d*_0_ is controlling the effect of the decay and is set to 5000. The set 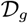 denotes the DHSs that are assigned to gene *g*. Details on the assignment are provided in the next section.

### Window based linkage

For each gene *g*, we consider a window *w* of size |*w*| centered at the most 5′-TSS of *g*. We denote all DNase peaks *d*, and the ChromHMM regions *m* that overlap *w* with 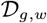 and 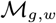, respectively. Thus, we set 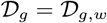 in equations 1–3 to compute scores based on DNase1-seq data.

Additionally, we define an intersection operation ∩_*H*_ between 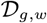and 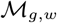 such that only 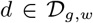 are retained that overlap by at least one 1*bp* with any 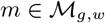. Formally, that is:

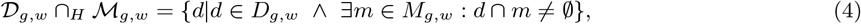

where *d* ∩ *m* indicates the overlap in genomic space of peak *d* and segment *m*.

We apply the ∩_*H*_ intersection operation to 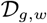 thereby obtaining 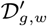:

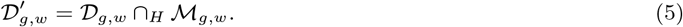

Consequently, we use 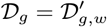 in equations 1–3 and compute scores as described above. The window based annotation is depicted in the upper part of Fig. 1.

### Nearest gene linkage

In this linkage paradigm, a peak (*d*) or segment (*m*) is exclusively associated to its closest gene. Notably this implies that a peak or segment can not be associated to more than one gene. Following this paradigm, we obtain 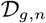, 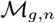 and set 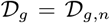 in equations (1)–(3). As above, in equations 4–5, we intersect 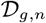 with 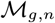 using the ∩_*H*_ operator and obtain 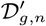. The nearest gene annotation is visualized in the middle of Fig.1.

### HiC and HiChIP based annotation

In addition to the window *w* centered at the TSS of gene *g*, we apply separate windows 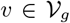 inferred from contacts of HiC or HiChIP experiments (equations 6–8). The set 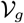 refers to all distant regions considered for gene *g*. We associate a chromatin contact to gene *g* if one of the two loop regions is located within a promoter search window of size *r*bp around the TSS of gene *g*. We refer to this search window as *loop window* (LW). The set of all DHSs intersecting a window 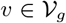 is denoted with 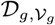, and are included in the score computation. Because the chromatin conformation capture experiment suggests a direct interaction of a potentially far away region *v* with gene *g*, we do not apply an exponential decay to peak signals of that region. However, we did test whether applying the exponential decay in the distal regions would be beneficial for model performance and found that it is indeed not the case for both HiC and HiChIP experiments, since all features were shrank towards zero (data not shown). Note that, in contrast to the promoter centric window *w*, there might be more than one window *v* for a distinct gene *g*.

In addition to the promoter centric features, we compute peak length *pl*_*g*_*, peak count *pc*_*g*_*, and peak signal *ps*_*g*_* in distal DHSs linked to gene *g* according to:

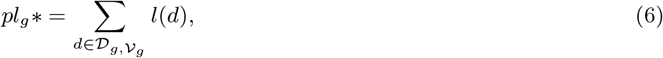

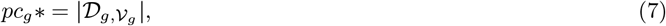

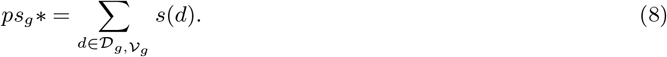

The HiC/HiChIP based annotation is explained in the bottom part of Fig. 1.

Finally, we integrate the ChromHMM information with the window based annotation. To this end, we intersect 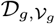 with 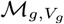 and obtain 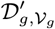 to reduce the number of regions associated to *g* from the distal regions 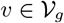 (Sup. Fig. 1).

### Computation of TF-gene scores with TEPIC

In addition to the peak based features *pl*_*g*_, *pc*_*g*_, *ps*_*g*_, 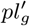, 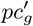, 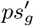 we estimate TF binding affinities using Tepic [24]. As introduced previously in Schmidt *et al.* [33], we compute TF affinities *a*_*p*,*t*_ for TF *t* in peak *p* using TRAP and aggregate the TF affinities to TF-gene scores *a*_*g*,*t*_ according to

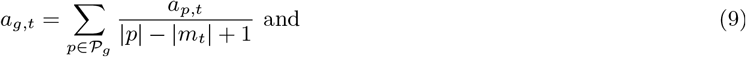

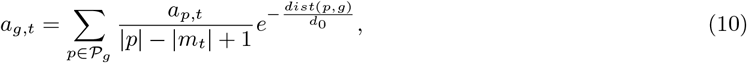

where 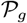 is the set of all DHSs assigned to gene *g*, reflecting the window based, or nearest gene assignment. The variable |*p*| denotes the length of DHS *p*, |*m*_*t*_| denotes the length of the Position Specific Energy Matrix (PSEM) *m*_*t*_ representing the binding preference of TF *t*, *dist*(*p*, *g*) is the distance between peak *p* and gene *g*, and *d*_0_ is a constant set to 5*kb* [23]. Here, we used 726 PSEMs for *Homo sapiens*, obtained from JASPAR [37], HOCOMOCO [38] and the Kellis ENCODE motif database [39], which are included in the TEPIC 2.0 repository [40].

As a baseline to be used in this study, we consider two promoter centric windows to compute TF-gene scores: (1) TF-gene scores aggregating TF affinities in a promoter window of size 3*kb* eq. 9, (2) TF-gene scores aggregating TF affinities in an extended promoter window of size 50kb including the exponential decay formulation of eq. 10. To utilise the information offered by the chromatin conformation capture data, we additionally compute TF-gene scores *a*_*g*,*t*_* solely based on DHSs overlapping the LWs:

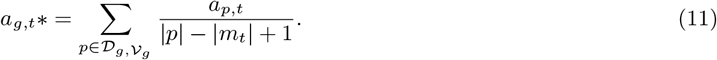

### Gene expression learning

Here, we briefly describe the machine learning techniques used in this study. An overview on the different feature setups is provided in Sup. Fig. 4, as well as in Table 1. The learning paradigm is sketched in Sup. Fig. 5.

**Table 1:** Different combinations of features evaluated in this study.

| Name | Considered Peak Features | Considered TF Features | Annotation |
| --- | --- | --- | --- |
| Promoter: Peaks | Peak Length $pl_g$<br>Peak Count $pc_g$<br>Peak Signal $ps_g$ | | Window<br>Nearest gene<br>ChromHMM |
| Promoter + HiC: Peaks<br>Promoter + HiChIP: Peaks | Peak Length $pl_g$<br>Peak Count $pc_g$<br>Peak Signal $ps_g$<br>Peak Length $pl_g^*$<br>Peak Count $pc_g^*$<br>Peak Signal $ps_g^*$ | | Window + HiC<br>Window + HiChIP<br>ChromHMM |
| Promoter + HiC: C Peaks<br>Promoter + HiChIP: C Peaks | Peak Length $pl_g + pl_g^*$<br>Peak Count $pc_g + pc_g^*$<br>Peak Signal $ps_g + ps_g^*$ | | Window + HiC<br>Window + HiChIP<br>ChromHMM |
| Promoter: Peaks + TFs | Peak Length $pl_g$<br>Peak Count $pc_g$<br>Peak Signal $ps_g$ | Affinities in promoter DHS $a_{g,t}$ | Window<br>ChromHMM |
| Promoter + HiChIP: EF Peaks + TFs | Peak Length $pl_g$<br>Peak Count $pc_g$<br>Peak Signal $ps_g$<br>Peak Length $pl_g^*$<br>Peak Count $pc_g^*$<br>Peak Signal $ps_g^*$ | Affinities in promoter DHS $a_{g,t}$<br>Affinities in loop DHS $a_{g,t^*}$ | Window + HiChIP<br>ChromHMM |

### Details on the linear model

Similar to a previous approach described in [24], we use linear regression with elastic net penalty implemented in the glmnet R-package [41] to predict gene expression. The elastic net combines two regularization terms, namely the Ridge (L2) and the Lasso (L1) penalty:

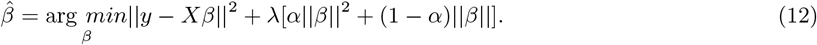

Here, the feature coefficient vector is represented by *β*, the estimated coefficients are denoted by 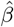, *X* refers to the feature matrix, *y* refers to the response vector, and the parameter λ determines the total amount of shrinkage. Both the input matrix *X* and the response vector *y*, containing gene expression estimates, are log-transformed, with a pseudo-count of 1, centered and normalized. The parameter *α*, which is optimized in a grid search from 0.0 to 1.0 with a step-size of 0.01, controls the trade-off between Ridge and Lasso penalty. Model performance is assessed on a hold-out test data set in a ten-fold outer Monte Carlo cross-validation procedure with 80% of the data randomly chosen to form the training data and 20% as test data. The λ parameter is fitted in a six-fold inner cross-validation using the *cv.glmnet* function. We choose the λ achieving the minimum cross validated error, computed as the average mean squared error (MSE) on the inner folds *(lambda.min)*.

### Details on the feature space

In this article, we build the feature matrix *X* in five different ways, listed in Table 1 and depicted in an exemplary manner in Sup. Fig. 4. As a baseline, we use our previously introduced promoter centric models considering DHS based features (*pl*_*g*_, *pc*_*g*_, *ps*_*g*_) and TF-gene scores *a*_*g*,*t*_. We refer to those as *Promoter: Peaks* and *Promoter: Peaks + TFs*, respectively.

Initially, we extended the promoter based models only with peak based features derived for loop sites (*pl*_*g*_*, *pc*_*g*_*, *ps*_*g*_*), due to simplicity. We refer to the separate consideration of promoter and loop peak features as *Promoter + HiC/HiChIP: Peaks* and to the combined consideration as *Promoter + HiC/HiChIP: C Peaks*.

Finally, we construct a feature matrix that is comprised of all peak (*pl*_*g*_, *pc*_*g*_, *ps*_*g*_, *pl*_*g*_*, *pc*_*g*_*, *ps*_*g*_*) and all TF features (*a*_*g*,*t*_, *a*_*g*,*t*_*). We refer to this distinct consideration of peak features and TF-gene scores for promoter DHS and enhancer DHS as the *extended feature space (EF)*, as it expands the original feature space considerably and allows a more detailed interpretation of the models.

### Gene sets

For all models in this study, protein coding genes (gencode v19) are considered.

### Implementation changes in TEPIC

We have extended the Tepic framework by a novel module that allows the integration of any matrix describing genome-wide chromatin contacts. The new module requires two inputs. Firstly, it requires a file with paired intervals (e.g. HiC or HiChIP loops) to be included in the annotation and secondly a parameter specifying the size of the loop window *LW* should be provided by the user. The *LW* is the area around a gene that is being screened for a potential chromatin contact. Accessible regions overlapping with a chromatin contact are not subject to the exponential decay. Furthermore, regions overlapping the promoter window as well as the *LW* are not counted twice. They are only considered for the promoter window to avoid redundancy. Details on the formatting of the required input file and on the novel parameters are provided in Sup. Sec. 3.

### TF gene expression analysis

We generated a mapping of TF names to Ensemble GeneIDs using Biomart. To test whether TFs in a query set have a higher expression than expected, we sampled 1000 TF sets of the size of the query set from the entire TF universe (without replacement). Whether the difference in the expression distributions is significant or not is assessed with a Wilcoxon test.

### Network analysis

Protein-protein interaction analysis was conducted using the STRING database version 11 [42]. We obtained the consensus PPI network for TFs (seed nodes) found in the extended feature space analysis (YY1, TCF7L2, TFDP1, REST, E2F8, E2F4, HOXA5, TEAD2, NRF1, TAL1, NR2F1, E2F4, EGR4). The final network was obtained using an interaction confidence score of 0.4 (default) and by allowing not more than five additional interactions with respect to the seed proteins (visualized in Fig. 6).

## Results

In this work, we developed an extension of our TEPIC approach that aggregates regulatory events occurring in potentially distal regulatory sites to the gene-level in a genome-wide fashion. Before we present the applicability and performance of this extension, we investigate common approaches that are widely applied in the community to establish PEIs add-hoc and use this comparison as a baseline for our novel methodology. Furthermore, we briefly describe differences in the HiC and HiChIP data used in this study.

### Local genomic architecture governs superiority of window or nearest gene based approaches

Previously, we have focused on window based linkage approaches in Tepic [24, 33]. Here, we have taken a broader scope and included the nearest gene assignment as well. Just like the window based approaches, this is another common strategy, used for instance by Gonzales *et al.* [25]. First we wanted to obtain a baseline, against which models considering chromatin conformation data can be compared later. We contrasted the performance of linear regression models predicting gene expression from the DNase1-seq derived features: (i) peak length, (ii) peak count, and (iii) peak signal for window based 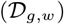 and nearest gene linkage 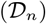 for GM12878, HeLa, HUVEC, IMR90, and K562 cells. As shown in Fig. 2 and in Sup.Fig. 2a, 50kb windows outperform 3kb windows and compared to nearest gene approaches, the 50kb window leads to slightly better models for three out of five samples. In Sup. Fig. 2b the mean squared error (MSE) for 9000 randomly selected, individual genes is shown using the DNase1-seq model for HeLa. Contrasting gene-specific prediction errors allows us to illustrate by comprehensive case-examples the existence of genome architecture specific advantages and disadvantages of the PEI linkage approaches.

**Figure 2:**
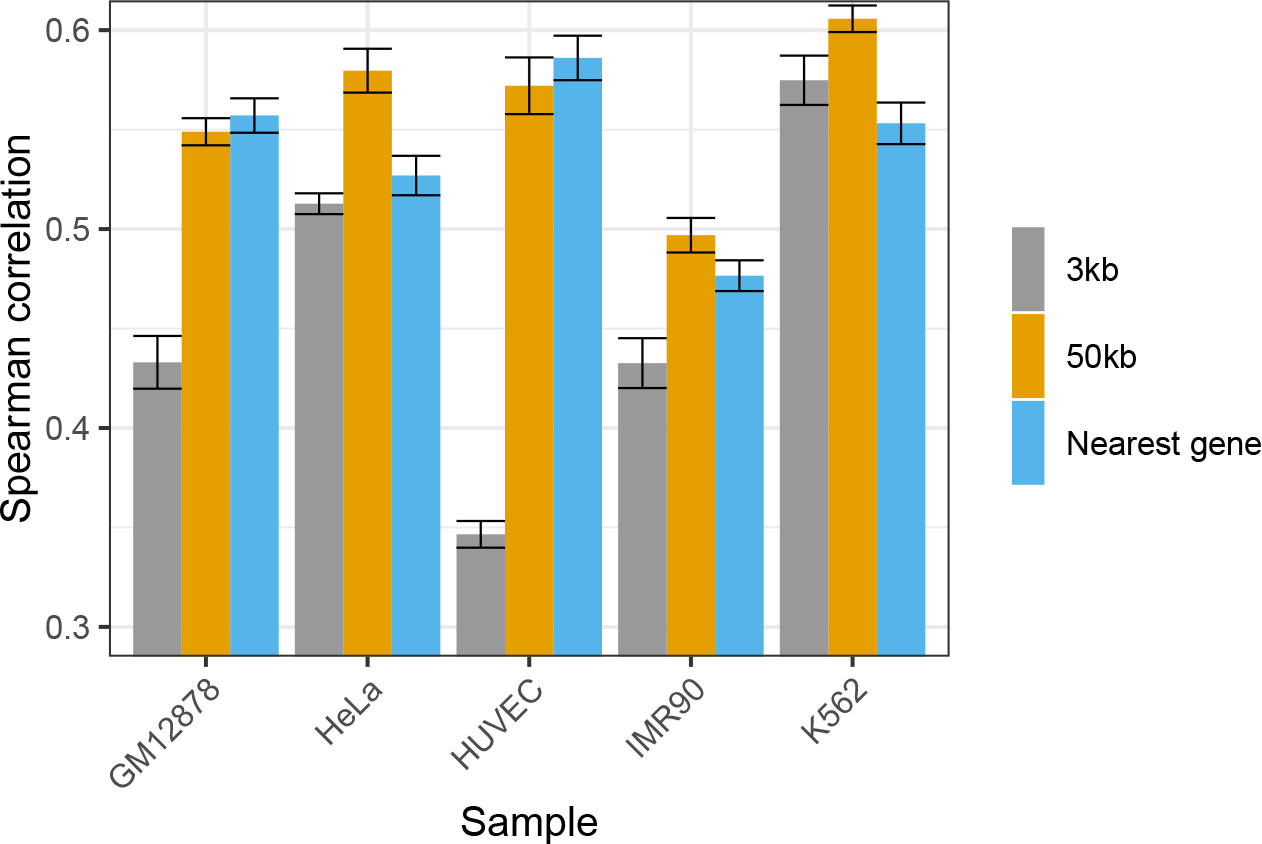
The performance of gene expression prediction models measured in terms of Spearman correlation on hold-out test data is shown for various models using peak length, peak count, and peak signal within the gene promoter regions. Two different window sizes (3kb, 50kb), and the nearest gene approach are compared. We observe that the 50kb models outperform the 3kb models. Considering the 50kb models, there is a slight advantage of the window based models over the nearest gene based annotation.

For example, the MSE of *RPL7A(ENSG00000148303)* is nearly twice as high using the nearest gene than the window based annotation. As shown in Sup. Fig. 3 there seems to be a bidirectional promoter for *RPL7A* and *MED22*. The model suggests that this can not be adequately covered by the nearest gene approach. A different scenario is depicted in Sup. Fig. 3b for the gene *HINT1(ENSG00000169567)*. This gene is located in a gene sparse region surrounded by several DHS peaks, which seem to add a large portion of noise in the nearest gene approach. In contrast to that, for the gene *APOA2(ENSG00000131096)*, the nearest gene approach leads to a better performance as it neglects, in contrast to the window based model, several DHS sites that seem to be associated to *TOMM40L* instead of *APOA2* (Sup. Fig. 3c). Each of these genes, *RPL7A, HINT1*, and *APOA2* is highlighted in Sup. Fig. 2b. Overall, these results suggest that neither the window based, nor the nearest gene annotation, generalise well across all genes. Still, the 50*kb* window based approach tends to perform slightly better on average. Therefore, we decided to augment the window based annotation using chromatin conformation capture data. Specifically, we attempt to replace the 50*kb* models with 3*kb* models that additionally consider DHSs linked to a distinct gene by chromatin conformation capture data. Only for reasons of comparison and to better understand the data at hand, we also augment the 50*kb* models with chromatin conformation data, although we note that in principle regions looped to the gene could be discovered using a chromatin conformation capture method.

### Including ChromHMM states is not beneficial for promoter-centric models

To understand whether the performance of the models could be improved by a stricter selection of potential regulatory regions, we used Promoter/Enhancer states predicted with ChromHMM in GM12878, HeLa, HUVEC, IMR90, and K562, thereby reducing the set of considered DHSs 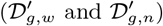. In Sup. Fig. 6 the results for the window and nearest gene annotation are contrasted. In general, the intersection of regulatory segments with DHSs 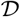 reduces model performance compared to the 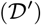 models. Only in case of HeLa, the nearest gene model does not loose performance.

This reduction in performance suggests that also relevant DHSs are removed from consideration. To investigate this hypothesis, we compared the mean q-value of removed DHSs in the 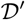 models compared to the retained DHS (Sup. Fig. 7a). Contrasting this hypothesis, we found that the q-value for the retained peaks is higher for both window sizes and the nearest gene approach. Note that in case of the nearest gene annotation, the ChromHMM intersection represents a genome-wide filtering. Also, a large portion of removed peaks are linked to *Quiescent/Low, Weak Repressed Polycomb* and *Weak transcription* chromatin states (Sup. Fig. 7b), which does not suggest that the removed regions have a regulatory role. Additionally, we observe that the average length of the considered DHSs tends to be shorter in 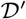 compared to 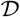 models (Sup. Fig. 8).

### HiC resolution impacts the association of genes to long range chromatin interactions

Before learning models using HiC or HiChIP data, we performed a few statistical analyses to better understand the characteristics of the chromatin conformation data. First, we assessed the overlap between DHSs and HiC as well as HiChIP loops, respectively. As shown in Fig. 3a, the fraction of HiC loops overlapping with at least one DHS increases with a decreasing HiC resolution. The tremendous differences between the various resolutions suggest that the choice of the used HiC resolution will likely affect any downstream analysis relying on DHS sites. Taking into account each HiC loop across all resolutions, at least 80% of the identified HiC loops intersect with at least one DHS site in four out of five cell lines. Compared to HiChIP data, sketched in Fig. 3b, it is striking how many genome-wide interactions are determined, compared to HiC data. For instance, in case of K562, there are 10, 000, 000 chromatin interactions with a DHS in both loop sites, contrasted against 6000 sites deduced from HiC data. As shown in Sup. Fig. 9a, there are still several magnitudes more HiChIP than HiC interactions, if a reduced HiChIP data set, filtered by q-Value or PET thresholding, is considered.

**Figure 3:**
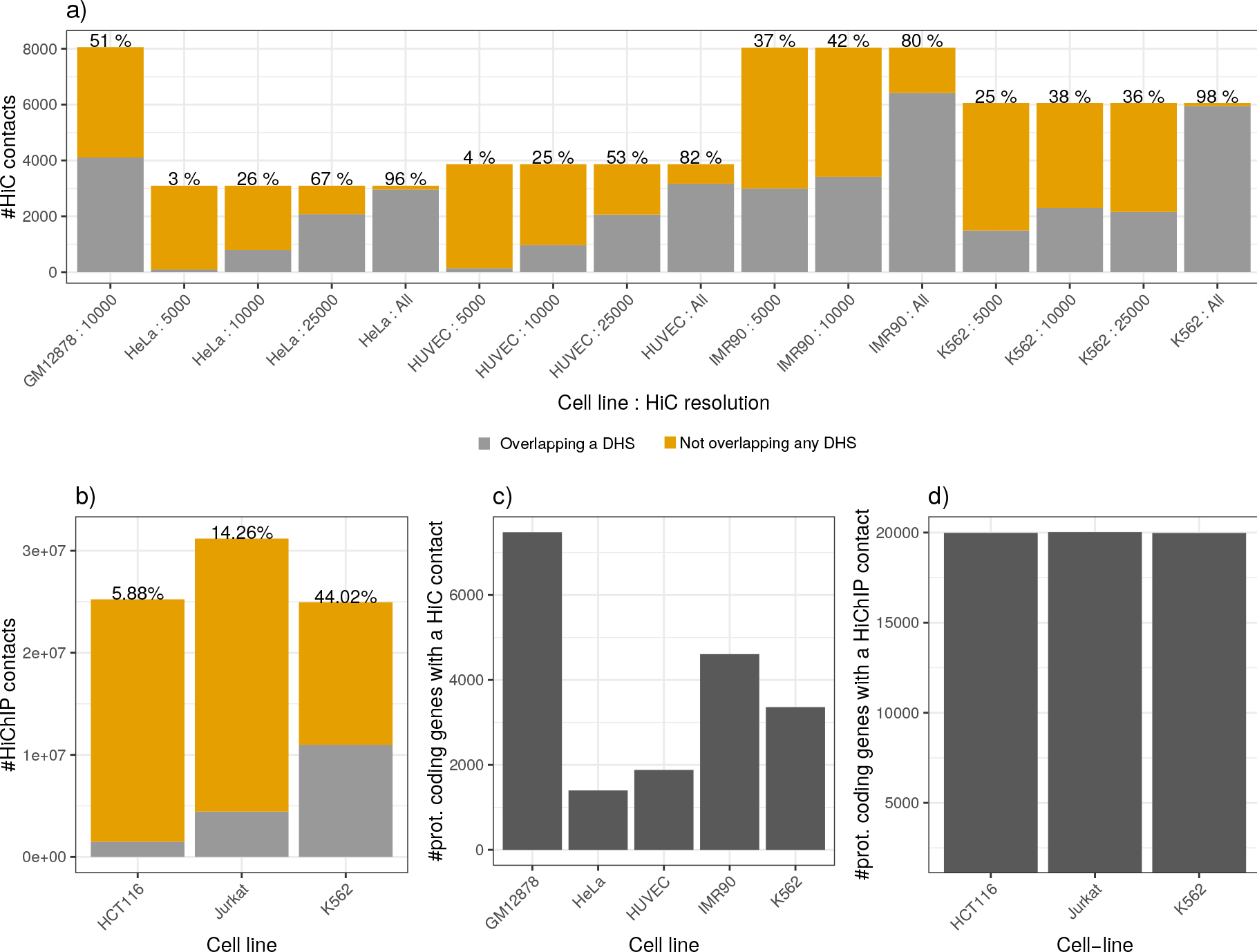
a) The number of HiC loops overlapping/not overlapping at least one DHS in each loop site is shown for different cell lines and different HiC resolutions. b) Analogously to a) for HiChIP contacts. All HiChIP samples have a resolution of 5*kb*. In a) and b), the ratio indicates the number of contacts overlapping DHS contrasted to those which do not. c) The bar plot shows the number of prot.coding genes that overlap a HiC loop using a *LW* of 25*kb* and a HiC resolution of 10*kb*. d) Analogously to c) but with HiChIP data and a promoter search window of 5*kb*.

As expected, the HiC resolution effects the number of genes that are linked to a chromatin loop. As exemplary shown in Fig. 3c for a *LW* of 25*kb* and a HiC resolution of 10*kb*, there are between 1000-7000 genes associated to a chromatin contact. In Sup. Fig. 9, we depict additional combinations for search windows and HiC resolutions. Generally, we observe that the number of genes associated to a loop reduces with a more precise, i.e. numerically smaller, HiC resolution. The *LW* used to link a HiC loop to a gene also influences the number of mapped genes. As expected, with an increasing search window size around the gene promoters, the number of genes that are linked to a loop is rising accordingly. Simultaneously the slope of the increase depends on the utilised HiC resolution. For example, as shown in Sup. Fig. 9b and c, the increase in the number of genes is only marginal for the best resolution (5*kb*), while it is more than three times as strong for the lowest one (25*kb*). With HiChIP data, almost all protein coding genes following the hg19 reference annotation are associated to a HiChIP contact (Fig. 3d). Upon a reduction of HiChIP contacts to those with a higher confidence, the number of affected genes stays above the levels of the HiC data (Sup.Fig.8b). As one might expect, the mean distance between HiChIP sites of one chromatin contact is decreasing with a more stringent thresholding (Sup. Fig. 9c). Together with the count information from Sup. Fig. 9a, we note that the HiChIP data contains between 10, 000-100, 000 interactions within a distance of 5*kb*-10*kb* between the interacting sites.

### Considering HiC and HiChIP data improves model performance

In addition to the promoter centric models shown in Fig. 2, we trained linear models additionally considering peak length, peak count, and peak signal of DHSs overlapping HiC and HiChIP loci, respectively. We refer to those features as *loop features*. Figure 4 illustrates that including HiC and HiChIP data is generally beneficial for model performance. For HiC data, depicted in Fig. 4a, we observe a slight improvement in model performance, which is more pronounced in case of a 3*kb* promoter window than with a 50*kb* promoter window. As observed in our earlier studies [33], the 50*kb* models outperform 3*kb* models. Overall, our results indicate that a larger LW tends to be beneficial for model performance. This is especially pronounced for GM12878, HUVEC, and IMR90 using a 3*kb* promoter window. This observation is likely to be directly linked to the dependence between the loop window size *LW* and the number of genes assigned to a HiC contact.

**Figure 4:**
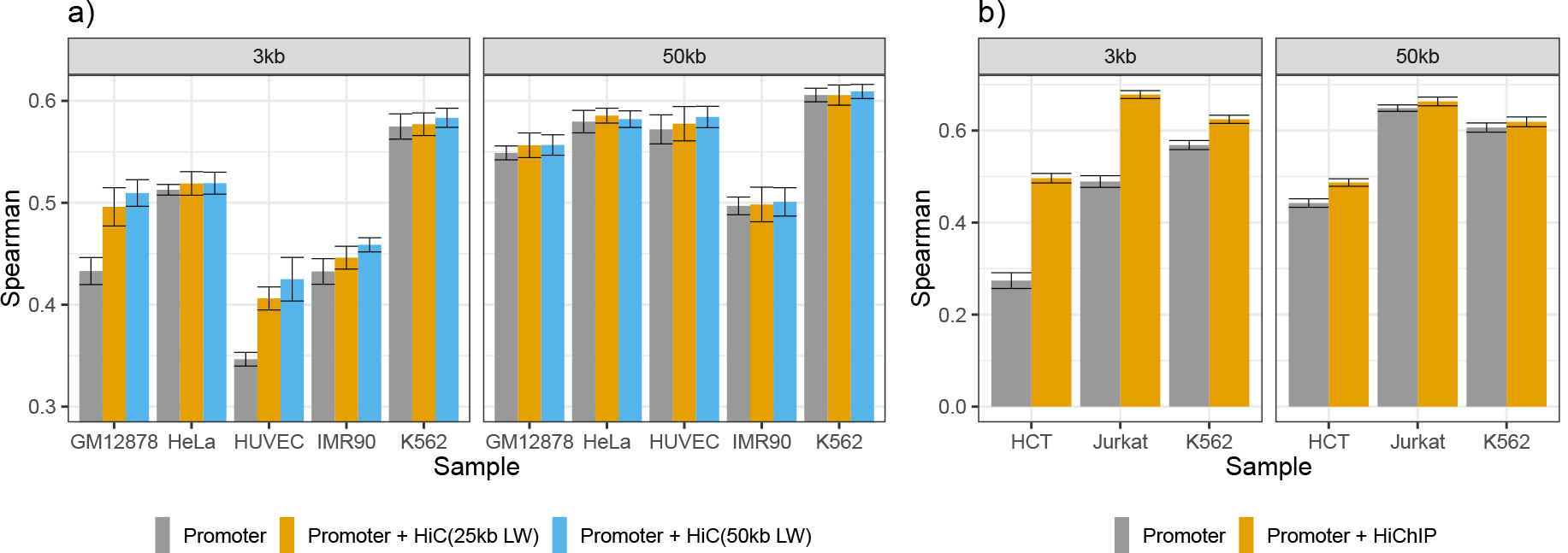
Model performance measured in Spearman correlation on hold-out test data considering chromatin contacts derived from a) HiC data using two different search window (25*kb* and 50*kb*) as well as b) using HiChIP data. We considered two different promoter windows, a 3*kb* and a 50*kb* window. While for HiC, the 50*kb* promoter windows led to better models than the 3*kb* models, it is the other way around considering HiChIP data.

In case of HiChIP data, illustrated in Fig. 4b, we see a stronger improvement of model performance upon inclusion of the loop features. Here, models extending the 3*kb* promoter window perform at least as good, or better than those extending the 50*kb* promoter window. It is possible, that the higher number of relatively short HiChIP interactions is responsible for this observation, which is in contrast to what we have seen with HiC data (Sup. Fig. 10a,c). As shown in Sup. Fig. 11, only a separate consideration of promoter and *loop features* leads to an improvement in model performance. Compared to promoter only models, filtering DHS with ChromHMM does improve models that are considering loop features (Sup. Fig.12).

We chose the 3*kb* HiChIP annotation for further examination in an extended feature space approach using TF affinities, because these models achieved the best performance with purely peak based features. As described in the next section, we attempt to decipher the regulatory impact of TFs binding in promoters and enhancers suggested by the chromatin conformation capture data.

### Results on the extended feature space

In our earlier works, we showed that models including TF affinities can be used to learn about the tissue specific regulatory activity of TFs [24, 33] and can be extended to investigate regulation of differential gene expression for example in Durek *et al.* [43]. Therefore we asked whether adding features for each TF would change the prediction performance. As shown in Fig. 5a, including TF affinities derived for DHSs around the promoter of genes, improves the performance of the linear models, compared to those models that are based solely on peak features.

**Figure 5:**
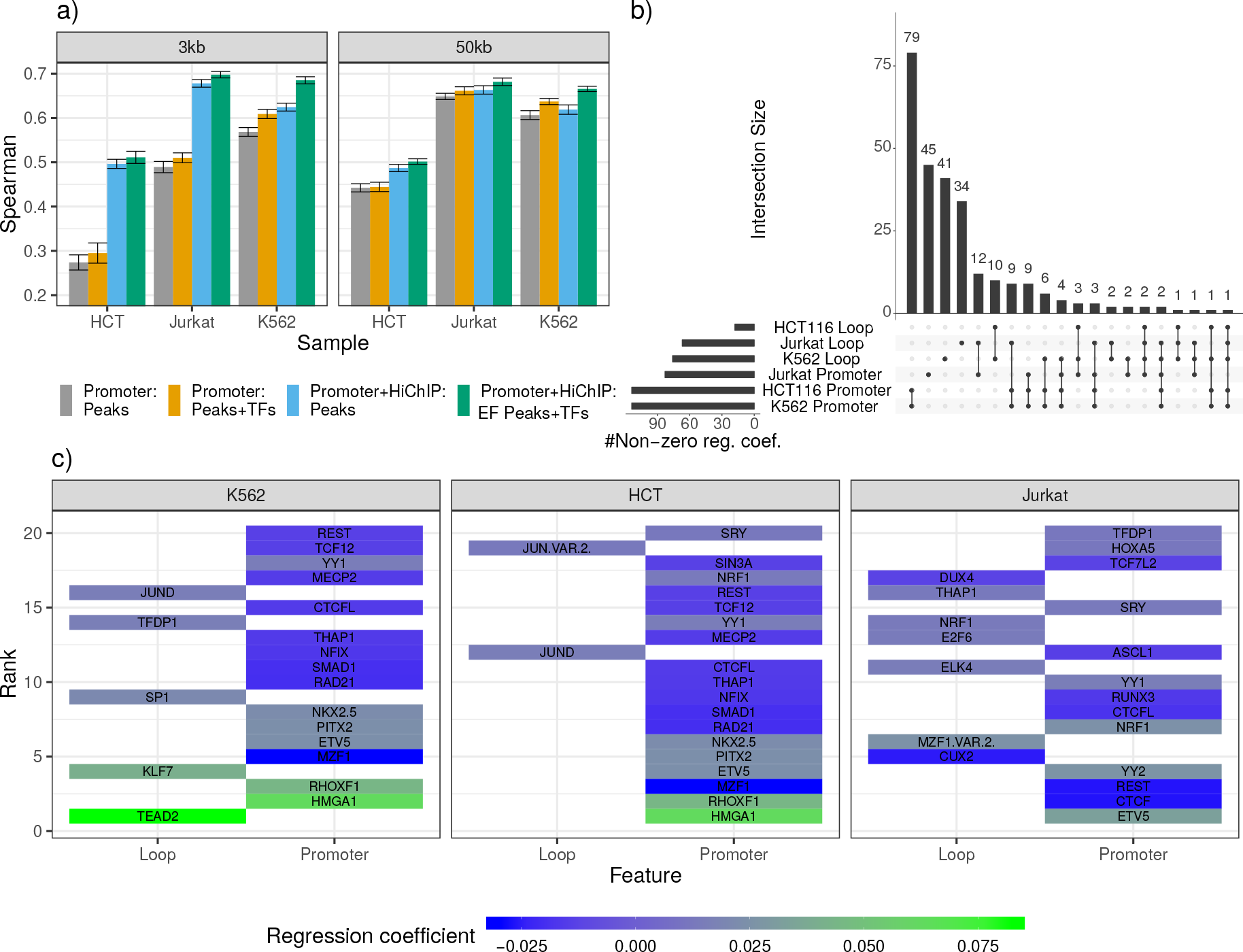
a) Model performance assessed in terms of Spearman correlation on hold out test data for models including TF-gene scores computed in the promoter and in the distal enhancers. Generally, including TF predictions improves model performance compared to considering only peak features. b) UpSet plot showing the relationship between TFs with a non-zero regression coefficients inferred by the extended feature space models. c) Ranking of the top 20 TFs by their absolute regression coefficients for each cell line. The color code indicates the mean regression-coefficient of the TFs computed in a 10-fold outer-cross validation.

Having the information about chromatin loops, we cannot only consider TF affinities in the promoter, but also in distal sites determined with HiChIP data. We trained models using an extended feature space considering each TF separately for the promoter and aggregated over the distal loop windows, for the K562, HCT116, and the Jurkat cell line. The distinct inclusion of these features further improves model performance (Fig. 5a, Sup. Fig. 13), suggesting that we can gain additional insights on the role of TFs by examining their regression coefficients. The UpSet plot in Fig. 5b depicts the overlap between TFs that have been assigned a non-zero mean regression coefficient in a 10-fold outer cross-validation procedure. The figure highlights that there are several factors occurring exclusively in promoter or loop regions, respectively.

Overall, we find the TFs BHLHE4, CTCFL, E2F8, ETV5, ETV6, HOXA5, NKX2.8, NKX3.1, NRF1, SRF, SRY, REST, RUNX1, TCF7L2, TEAD2, TFDP1, YY1, and YY2 to be commonly selected as a feature in the promoter region for all three cell lines. The TFs E2F4, EGR4, NR2F1, and TAL1, are commonly selected in the loop regions.

Recalling that the HiChIP data was performed with an antibody targeting YY1 and the fact that YY1 binding sites are over represented in human core promoters [44], the prediction of YY1 as a common promoter feature is a validation of our computational approach. The appearance of YY2 may be due to the fact that the C-terminal binding domains of YY1 and YY2 are highly conserved. Indeed, ChIP-seq derived binding peaks of YY2 contained the YY1 motif at peak centers, indicating that YY2 binds similar regions and that there is an overlap of genes regulated by both TFs [45, 46].

Similar to YY1, also NRF1 has been shown to be essential for transcriptional regulation at the core promoter of several genes [47, 48]. The TF TFDP1 has been shown to bind to the promoter elements of genes that are related to the cell cyle [49]. Regarding the TFs commonly identified in loop regions, we find, for instance, that E2F4 has been suggested previously to bind to enhancer regions [50]. Binding sites of NR2F1 have been shown to coincide with high levels of the established enhancer marks P300 and H3K27ac [51].

Taking into account that the HiChIP data suggests a spatial proximity between the TFs bound to the promoter regions to those TFs bound to the loop regions, we investigated protein-protein interactions using the STRING database [42]. We selected all TFs that occur in our models in all three cell lines, in either promoter or loop regions. The resulting network is shown Fig. 6. We find that there are many confident interactions between several of the selected factors, including YY1, NRF1, HOXA5, EGR4, E2F4, TFDP1, TCF7L2, and TEAD2 suggesting the formation of protein-protein complexes ultimately linking enhancer and promoter regions.

In addition to this general analysis, we investigated the top 20 TFs ranked by their mean, absolute, regression coefficient across the 10-fold outer cross-validation, per cell line (Fig. 5c). A prominent example for a TF selected in the promoter region for K562 and HCT116 is HMGA1. This TF is known to act as an essential regulator for the mediator complex and the basal transcription machinery [52]. Also YY1 is among the top 20 TFs selected for each cell line, supporting the validity of the ranking. On the loop sites, we find for example JUND being among the top 20 factors in K562 and HCT116. This factor is known to support enhancer functions for instance in B cells and keratinocytes [53, 54]. Interestingly, the knockdown of SP1 or KLF7, which are selected for K562 cells among the top loop features, has been shown to impact cellular differentiation and *β*-globin production, respectively [55].

The potential regulatory role of the TFs highlighted in Figs. 5c and 6 is supported by the observation that the top 50 TFs with a non-zero regression coefficient per cell line have higher gene expression values than randomly sampled TFs (Sup. Fig. 14).

**Figure 6:**
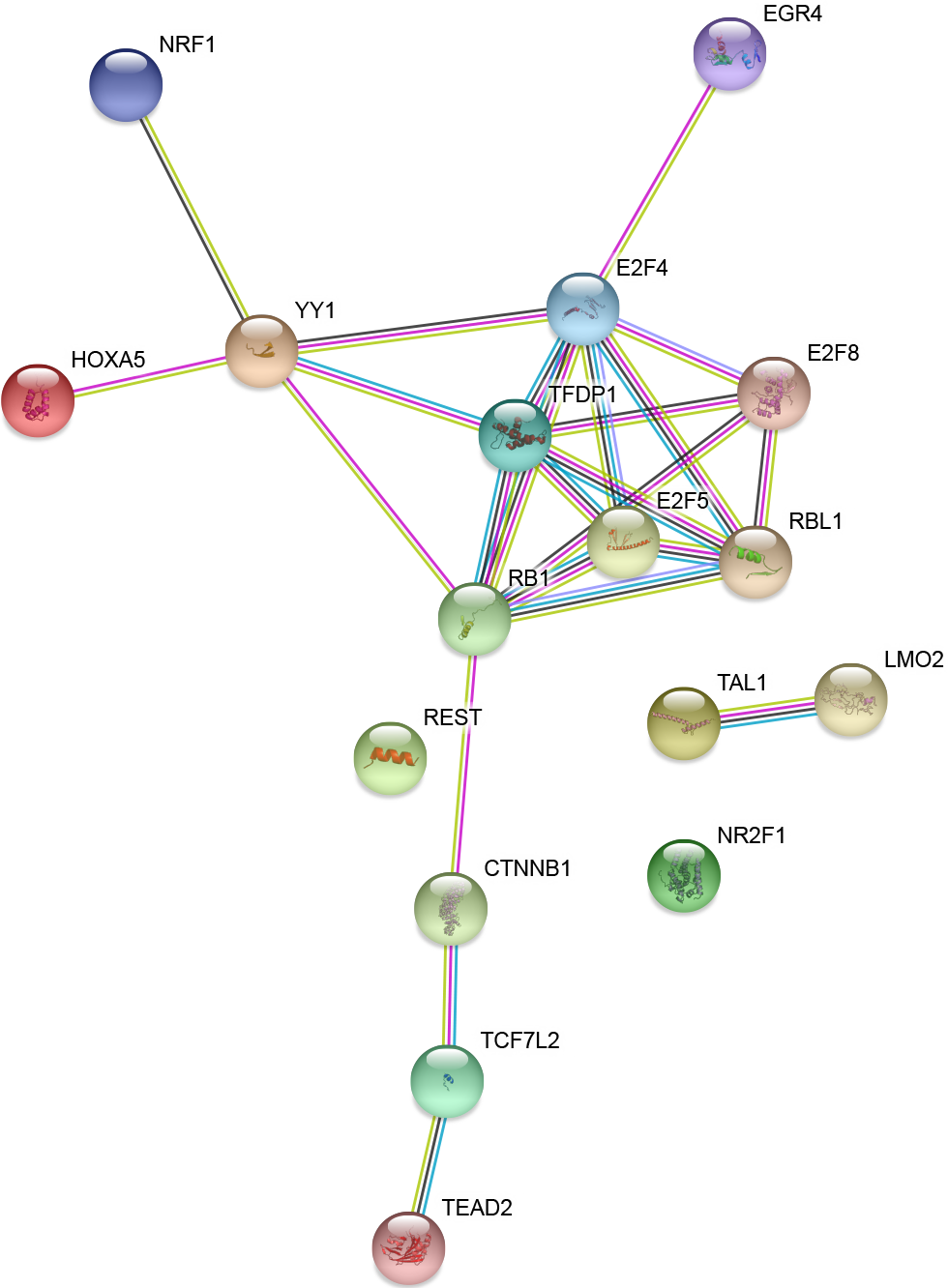
Protein-protein interaction network obtained from the STRING database illustrating interactions among YY1 and TFs commonly selected as a predictor in gene expression models for K562, Jurkat, and HCT116 in promoter and distal loop sites.

The analysis thus revealed a set of core TFs that are involved in enhancer-loop linkage involving the TF YY1.

## Discussion

Associating regulatory regions to genes is still subject to ongoing research. In this work, we compared established methods to construct PEIs and present an extension of our TEPIC approach to associate enhancers with genes using chromatin conformation capture data, exemplary using HiC and HiChIP data. We evaluated the different PEI linkage methods using predictive models of gene expression learned on DNase1-seq data.

Our results indicate that neither the widely used nearest-gene linkage nor the window based PEI models are optimal. We illustrate that both approaches have distinct advantages and drawbacks. For example, in the nearest gene assignment, there is no common agreement, whether the TSS or TTS of a gene is used to calculate the genes’ distance to the putative enhancer. Also, in gene dense regions, it is not obvious whether a peak should be uniquely assigned to only one or to multiple genes. Indeed, it was shown before that a distinct enhancer can influence the expression of various genes [56, 57]. On the other hand, the window based linkage might generate many false positive associations in gene-dense regions and likely misses distal enhancer regions, which in turn might be captured by the nearest gene approach in gene-sparse genomic loci. We have illustrated these points using multiple examples in Sup.Fig. 2. Notably, current research suggests that many enhancer-gene interactions are established only within TADs but only rarely across TAD borders [32]. This might argue in favour of window based approaches and suggests to include TAD boundaries in nearest gene approaches to avoid assignments across TAD boundaries. It is of crucial importance for the field to understand the pros and cons of the assignment strategies, because they still form the basis for recent efforts trying to link enhancers to genes in multi-tissue scenarios [27]. Also, in settings were only few samples are available, a computational *de novo* assignment of regulatory regions to genes using correlation based methods is not feasible.

Our examination of the available HiC data suggests that the peak resolution has a strong impact on inferring PEIs. We showed that both the number of genes as well as the number of overlapping DHS sites largely depends on the HiC resolution. Notably, with a higher, i.e. numerically smaller, HiC resolution the number of genes associated to HiC loops almost remains constant with an increasing search window size. While models considering HiC data did improve in their ability to predict gene expression, the improvement was only marginal compared to the 50kb promoter model. This can be due to several reasons. One possibility might be that not all chromatin contacts are directly linked to transcriptional regulation and gene expression, as also suggested by Ray et al. [58]. For instance, loops could be part of higher-order structures and thereby indirectly influence cellular processes. This could be an explanation for the varying overlap of DHSs with HiC loops described in Fig.5. It is likely that other methods, e.g. ChIA-PET, capture HiC[59], or HiChIP, which can enrich the sequencing libraries for distinct regions such as promoters of interest, lead to more precise contact maps in terms of both resolution and signal-to-noise ratio. Leveraging these more fine-grained technologies for gene expression modeling seems to be an opportunity to improve the prediction performance and to enhance our understanding on the underlying regulatory processes.

To examine this hypotheses, we learned gene expression prediction models using recently generated HiChIP data. In contrast to the HiC data, we see a stronger improvement in model performance. Importantly, the rather tight 3kb window focusing at the promoter augmented with HiChIP contacts, outperformed all 50kb model variants, suggesting that the interactions suggested by the HiChIP experiments are indeed meaningful and better than an average over all DHSs within 50kb of the TSS. As depicted in Fig. 4, the number of HiChIP contacts is several magnitudes higher than the number of HiC contacts. We noted that many of the high quality HiChIP contacts belong smaller range contacts, suggesting that HiChIP data also uncovers chromatin interactions between a gene’s TSS and intragenic enhancers. This might explain why the augmented 3kb models perform at least as good or better than the 50kb models as intragenic interactions are likely to be modelled in the HiChIP data.

In contrast to the promoter only models, including ChromHMM state segmentations did improve the performance of models considering HiC or HiChIP data, suggesting that both techniques lead to the inclusion of less relevant DHSs.

Wrapping up all these aspects, we tried to further improve the HiChIP models by incorporating TF affinities into the extended feature space. They allow the inferred regulators to be prioritized in a promoter and a (distal) enhancer specific point of view. We have shown that models based on this feature design can be comprehensively interpreted and lead to biologically meaningful and reasonable results.

To make our approach generally applicable for the research community and to scale-up with new experimental technologies, we designed TEPICS chromatin conformation extension to be able to integrate PEIs derived from any chromatin conformation capture technology. We believe that this extension together with the extended feature space annotation will be helpful to elucidate regulatory processes at promoters and enhancers.

### Conclusion

Overall, our study provides an unbiased comparison of prevalent PEI linkage strategies and shows that neither the established window based PEI linkage nor the nearest gene linkage perform optimal. Further, we show that HiC and HiChIP data can both be used to integrate genome-wide chromatin contacts into predictive gene expression models. Thereby, we can not only improve model performance, but, using our extended feature space formulation, enable users to obtain detailed insights into the promoter and enhancer specific activity of TFs across distinct cell types and tissues.

## Availability of data and materials

The code used to generate the results in this work is available online: https://github.com/schulzlab/TEPIC. Data identifiers are provided in Supplementary Tables 1,3, and 4.

## Author’s contributions

FK performed the HiC study and the window based versus nearest gene annotation comparison. FS generated the HiChIP models and performed related analyses. FS and MHS advised FK. FS wrote the manuscript and, together with FK, generated the figures. FK and MHS commented on the manuscript. MHS supervised the study.

## Acknowledgments

We thank the ENCODE consortium for providing and processing the RNA-seq and DNase1-seq data as well as the Lieberman-Aiden group for sharing their HiC datasets and downstream applications.

