## Supplementary Material for "Integrative prediction of gene expression with chromatin accessibility and conformation data"

#### 1. Supplementary Tables

| **ENCODE accession number** | **Data Type** |
|---|---|
| ENCFF000DYC | Quantified mRNA of K562 |
| ENCFF441RET | DNase -1 seq of K562 |
| ENCFF000CZF | Quantified mRNA of GM12878 |
| ENCFF000SKV | DNase -1 seq of GM12878 |
| ENCFF000DJU | Quantified mRNA of IMR90 |
| ENCFF000SOC | DNase-1 seq of IMR90 |
| ENCFF000DNW | Quantified mRNA of HeLa |
| ENCFF000SPR | DNase-1 seq of HeLa |
| ENCFF000DUQ | Quantified mRNA of HUVEC |
| ENCFF001DNS | DNase-1 seq of HUVEC |
| ENCFF673ODZ | Quantified mRNA of JURKAT |
| ENCFF164FDV, ENCFF813IXN | DNase1-seq of JURKAT |
| ENCSR000CWM | RNA-seq reads of HCT116 |
| ENCFF081DDV, ENCFF291HHS | DNase1-seq of HCT116 |
| ENCFF916QPX | ChromHMM states for K562 |
| ENCFF869GUF | ChromHMM states for GM12878 |
| ENCFF147PPH | ChromHMM states for IMR90 |
| ENCFF654HNG | ChromHMM states for HeLa |
| ENCFF3970PB | ChromHMM states for HUVEC |

**Supplementary Table1:** Identifiers of ENCODE RNA-seq, DNase-1-seq, TF-ChIP-seq data, and ChromHMM files.

| Cell line | Number of identified DHS |
|---|---|
| **K562** | 951.681 |
| **GM12878** | 77.863 |
| **IMR90** | 487.220 |
| **HeLa** | 673.093 |
| **HUVEC** | 164.551 |
| **HCT116** | 153.738 |
| **JURKAT** | 322.906 |

**Supplementary Table2:** Sample Identifiers and description

| Supplement Identifier | Cell-line | Available resolutions |
|---|---|---|
| **GSE63525_GM12878_primary_HiCCUPS_looplist.txt.gz** | GM12878 | 10kb |
| **GSE63525_HUVEC_HiCCUPS_looplist.txt.gz** | HUVEC | 5kb, 10kb, 25kb |
| **GSE63525_HeLa_HiCCUPS_looplist.txt.gz** | HeLa | 5kb, 10kb, 25kb |
| **GSE63525_IMR90_HiCCUPS_looplist.txt.gz** | IMR90 | 5kb, 10kb |
| **GSE63525_K562_HiCCUPS_looplist.txt.gz** | K562 | 5kb, 10kb, 25kb |

**Supplementary Table3:** Overview on the Hi-C data used in this study. The original data was obtained from Gene Expression Omnibus using the accession number GSE63525.

| Supplement Identifier | Cell-line |
|---|---|
| **GSE63525_GM12878_primary_HiCCUPS_looplist.txt.gz** | GM12878 |
| **GSE63525_HUVEC_HiCCUPS_looplist.txt.gz** | HUVEC |
| **GSE63525_HeLa_HiCCUPS_looplist.txt.gz** | HeLa |

**Supplementary Table4:** Overview on the Hi-ChIP data used in this study. The original data was obtained from Weihrauch et al.

#### 2. Supplementary Figures

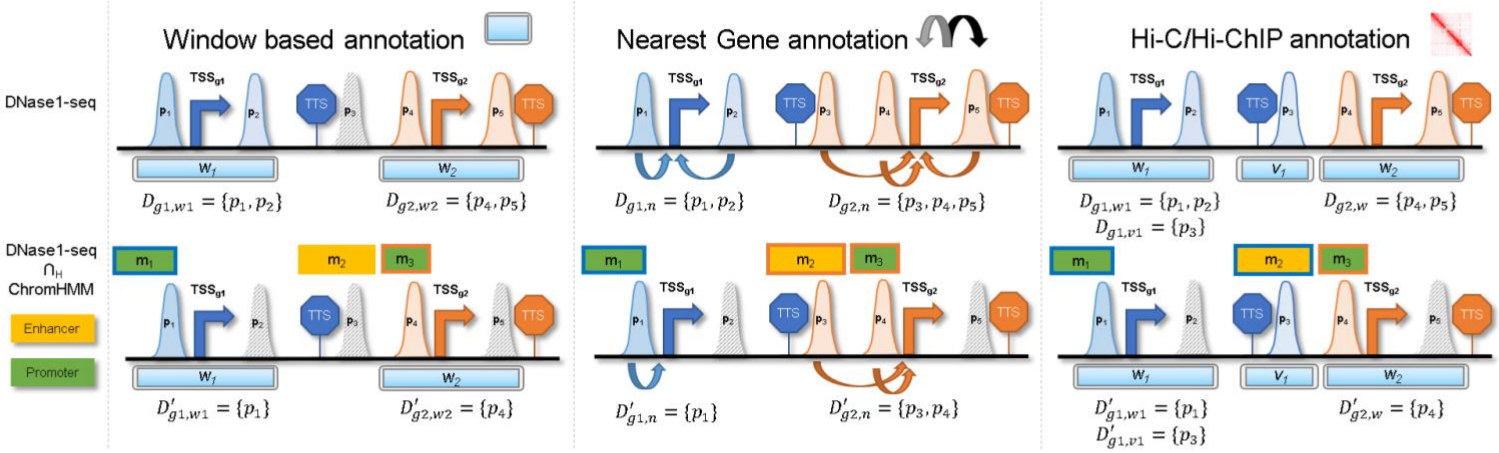


**Supplementary Figure 1:** Here, the ChromHMM based filtering of the annotation versions shown in Figure 1 is shown. Briefly, only DHSs overlapping a ChromHMM promoter or enhancer segment are considered.

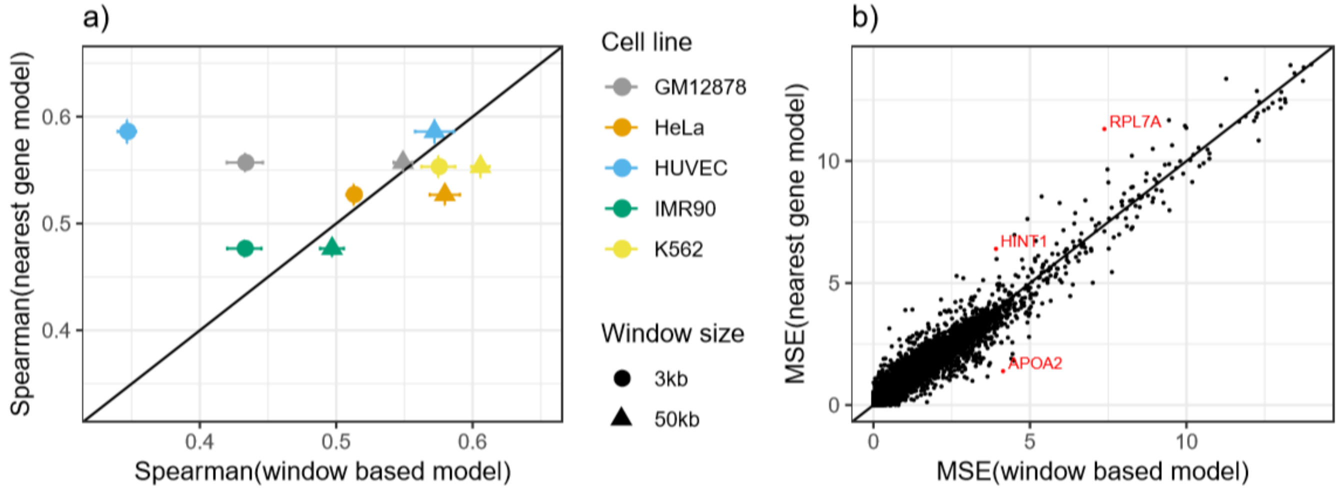


**Supplementary Figure 2:** a) Spearman correlation achieved by regression models of gene expression using aggregated DNase1-seq data for a window based (x-axis) and nearest gene based (y-axis) enhancer linkage. On average the window based approaches are outperforming the nearest gene association. In b) the mean squared error (MSE) between predicted and measured gene expression for $9000$ randomly selected HeLa genes is shown for both window based and nearest gene models. For genes highlighted in red, Sup. Fig. 2 shows IGV screen-shots illustrating the chromatin landscape around them.

a)

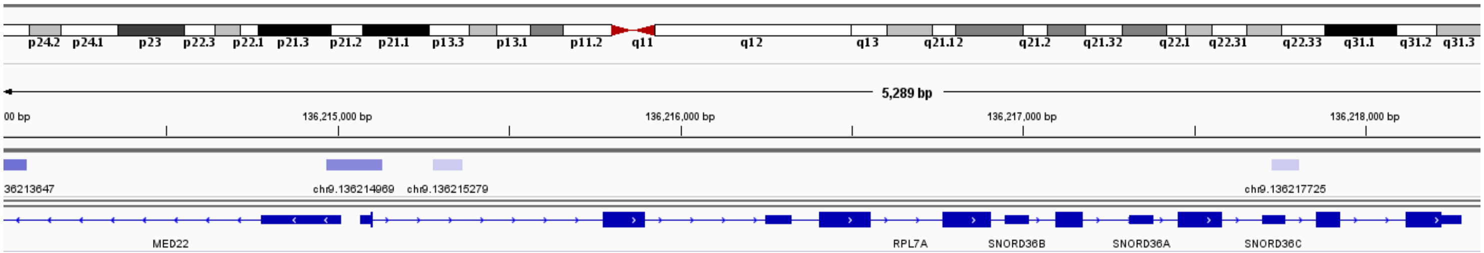


b)

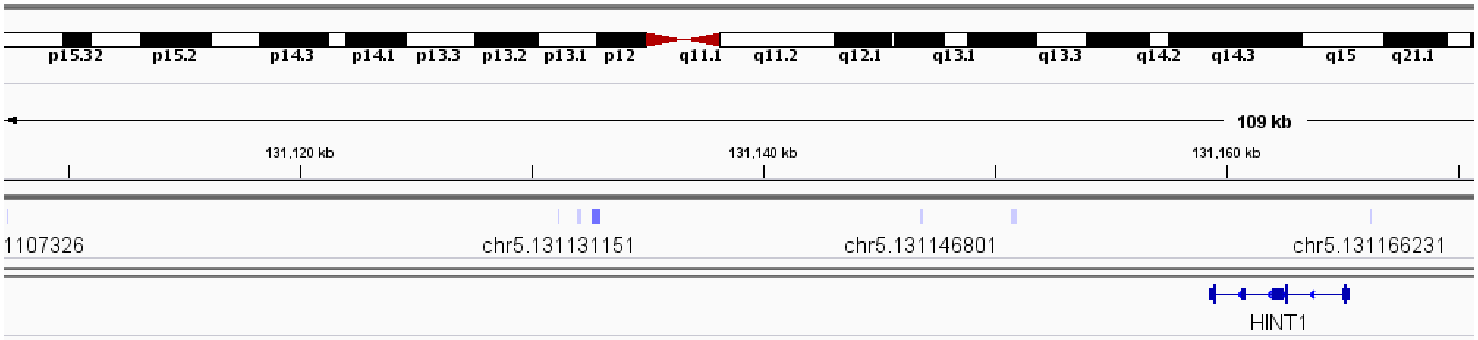


c)

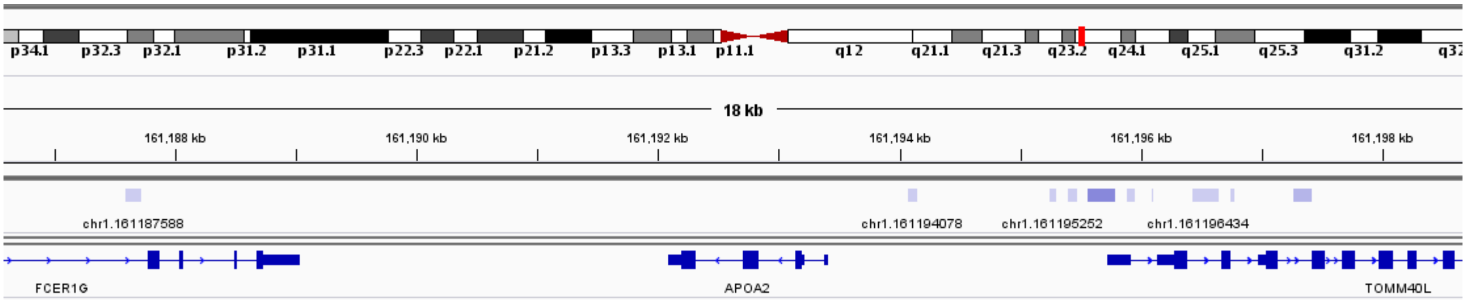


**Supplementary Figure 3:** IGV shots showing the JAMM peaks called in HeLa centered at three genes: a) RPL7A, b) HINT1, c) APOA2.

**a)** Promoter: Peaks

| | **Peak count** | **Peak length** | **Peak signal** |
|---|---|---|---|
| **Gene 1** | $pc_1$ | $pl_1$ | $ps_1$ |
| **…** | | | |
| **Gene m** | $pc_m$ | $pl_m$ | $ps_m$ |

**b)** Promoter + HiC/HiChIP: Peaks

| | **Peak count** | **Peak length** | **Peak signal** | **Peak count*** | **Peak length*** | **Peak signal*** |
|---|---|---|---|---|---|---|
| **Gene 1** | $pc_1$ | $pl_1$ | $ps_1$ | $pc_1{}^*$ | $pl_1{}^*$ | $ps_1{}^*$ |
| **…** | | | | | | |
| **Gene m** | $pc_m$ | $pl_m$ | $ps_m$ | $pc_m{}^*$ | $pl_m{}^*$ | $ps_m{}^*$ |

**c)** Promoter + HiC/HiChIP: C Peaks

| | **Peak count** | **Peak length** | **Peak signal** |
|---|---|---|---|
| **Gene 1** | $pc_1 + pc_1{}^*$ | $pl_1 + pl_1{}^*$ | $ps_1 + ps_1{}^*$ |
| **…** | | | |
| **Gene m** | $pc_m + pc_m{}^*$ | $pl_m + pl_m{}^*$ | $ps_m + ps_m{}^*$ |

**d)** Promoter: Peaks + TFs

| | **Peak count** | **Peak length** | **Peak signal** | **Affinities TF t** |
|---|---|---|---|---|
| **Gene 1** | $pc_1$ | $pl_1$ | $ps_1$ | $a_{1,t}$ |
| **…** | | | | |
| **Gene m** | $pc_m$ | $pl_m$ | $ps_m$ | $a_{m,t}$ |

**e)** Promoter + HiChIP: EF Peaks + TFs

| | **Peak count** | **Peak length** | **Peak signal** | **Peak count*** | **Peak length*** | **Peak signal*** | **Affinities TF t** | **Affinities* TF t** |
|---|---|---|---|---|---|---|---|---|
| **Gene 1** | $pc_1$ | $pl_1$ | $ps_1$ | $pc_1{}^*$ | $pl_1{}^*$ | $ps_1{}^*$ | $a_{1,t}$ | $a_{1,t}{}^*$ |
| **…** | | | | | | | | |
| **Gene m** | $pc_m$ | $pl_m$ | $ps_m$ | $pc_m{}^*$ | $pl_m{}^*$ | $ps_m{}^*$ | $a_{m,t}$ | $a_{m,t}{}^*$ |

**Supplementary Figure 4:** The different feature matrices used in this study are shown.

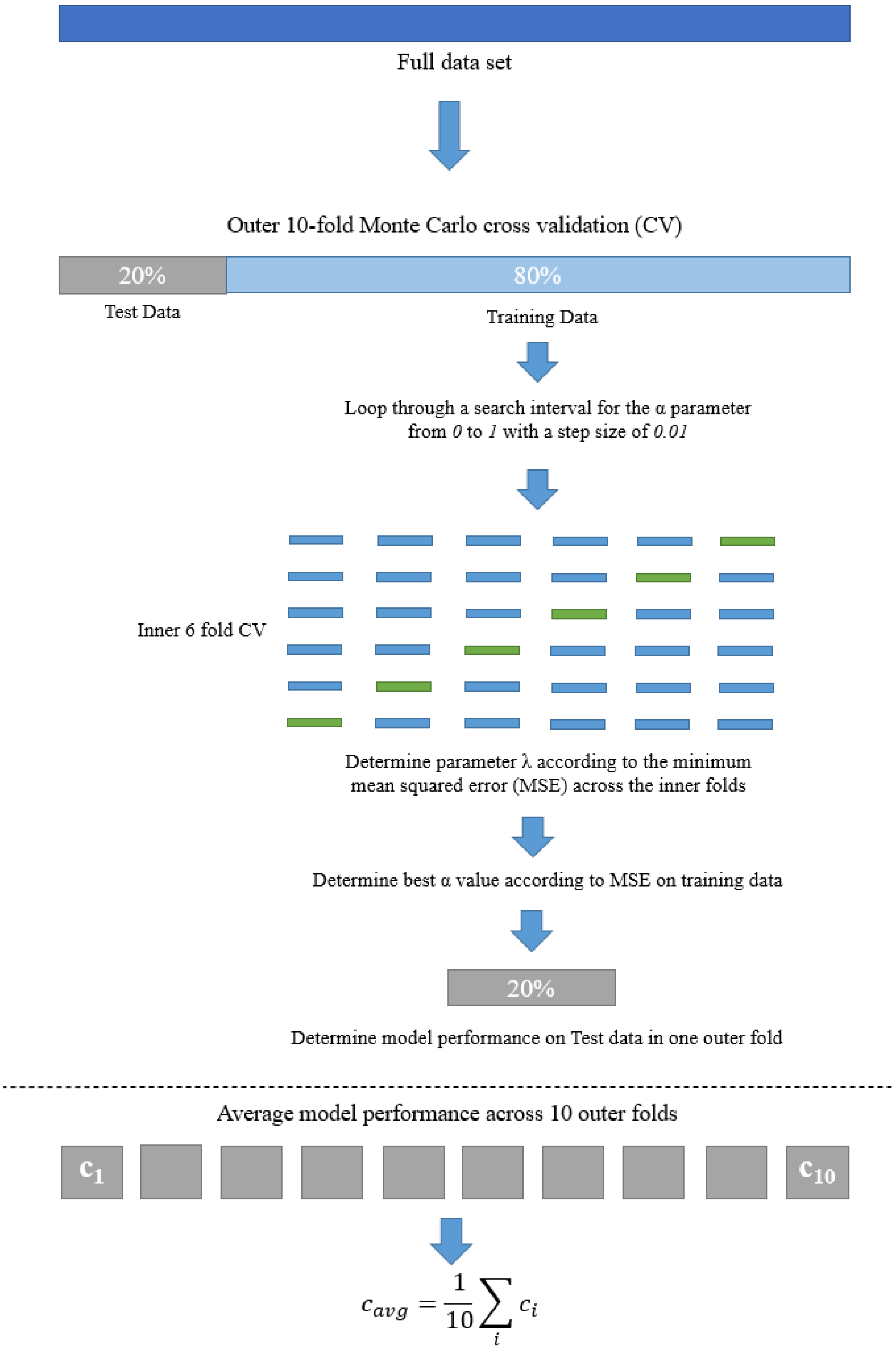


**Supplementary Figure 5:** A scheme of the used linear regression paradigm.

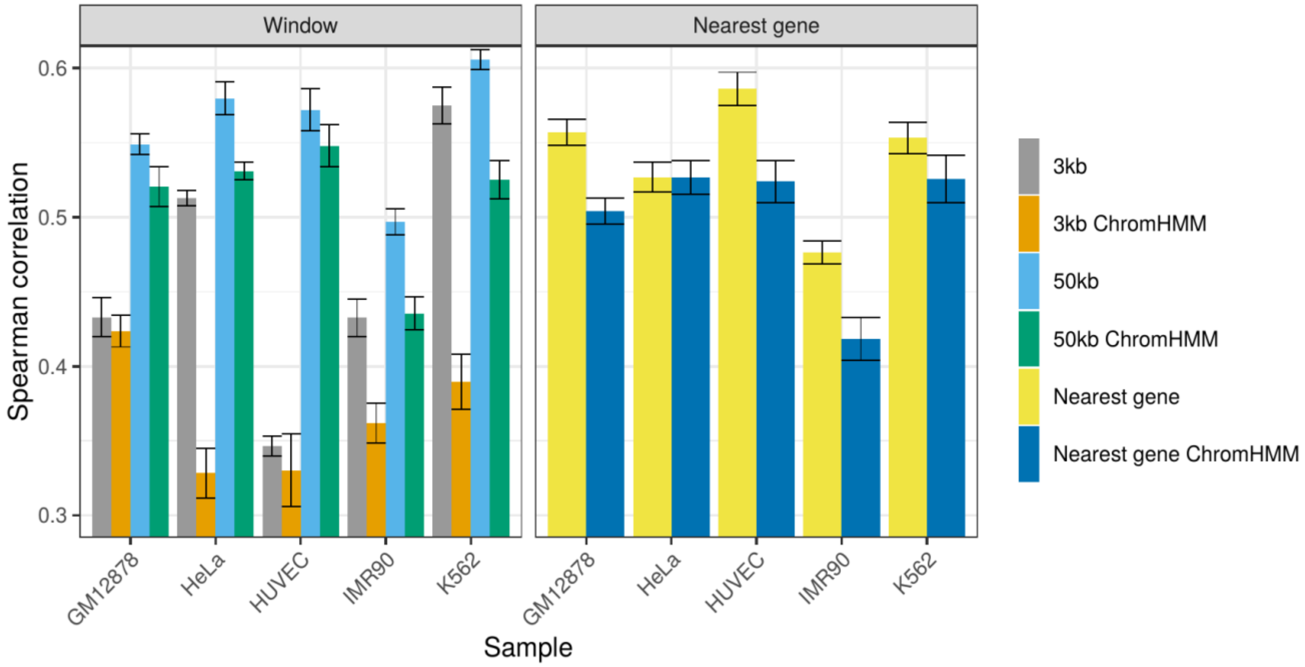


**Supplementary Figure 6**: Here, model performance is shown in terms of Spearman correlation for Window and nearest gene based approaches with and without an additional filtering with ChromHMM.

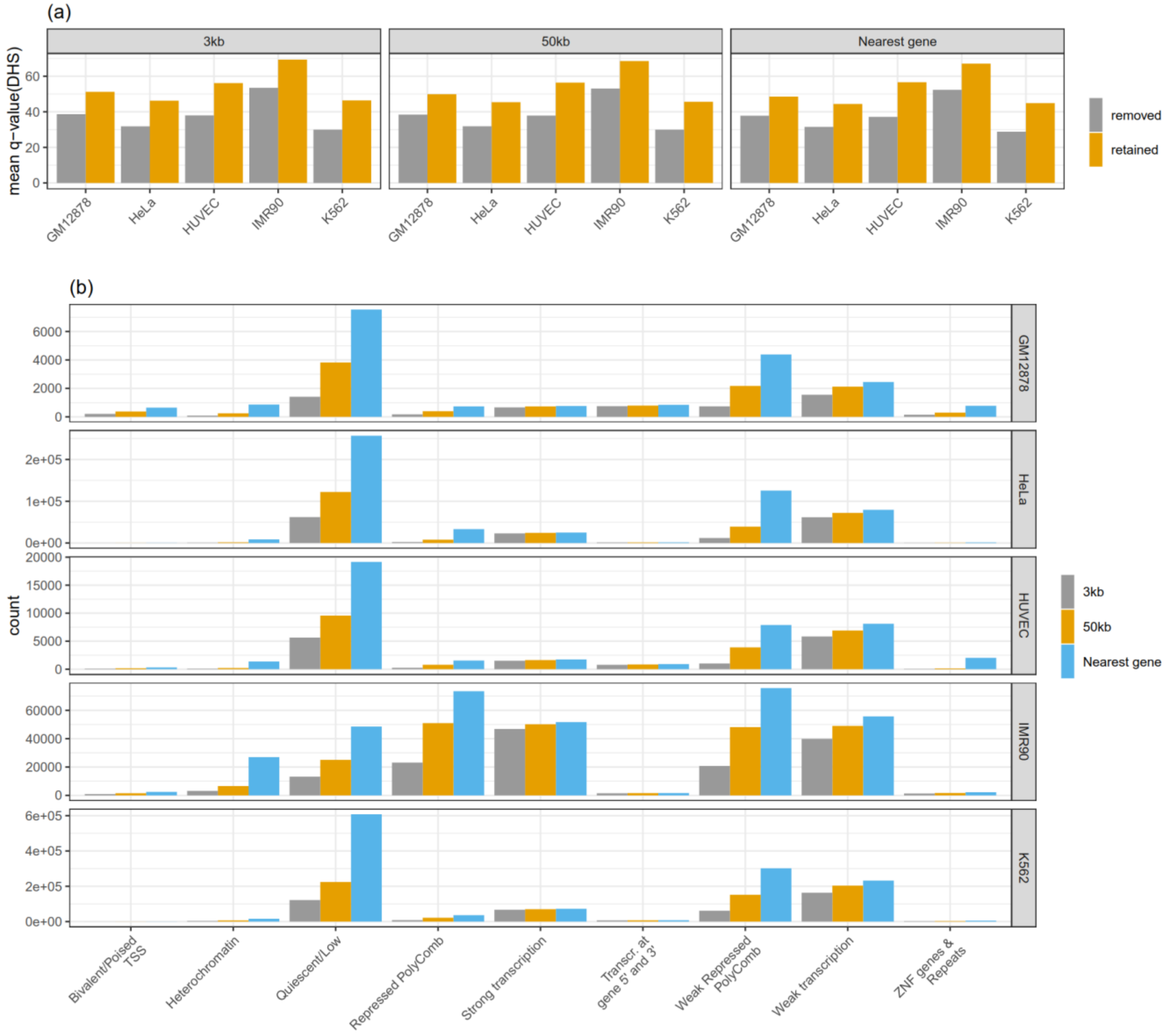


**Supplementary Figure 7:** a) shows details on the characteristics of DNase1-seq peaks retained/removed by intersection with ChromHMM Promoter/Enhancer states. In part (a) the score of retained/removed peaks is shown, in (b) the count of removed peaks is shown per overlapping chromatin state.

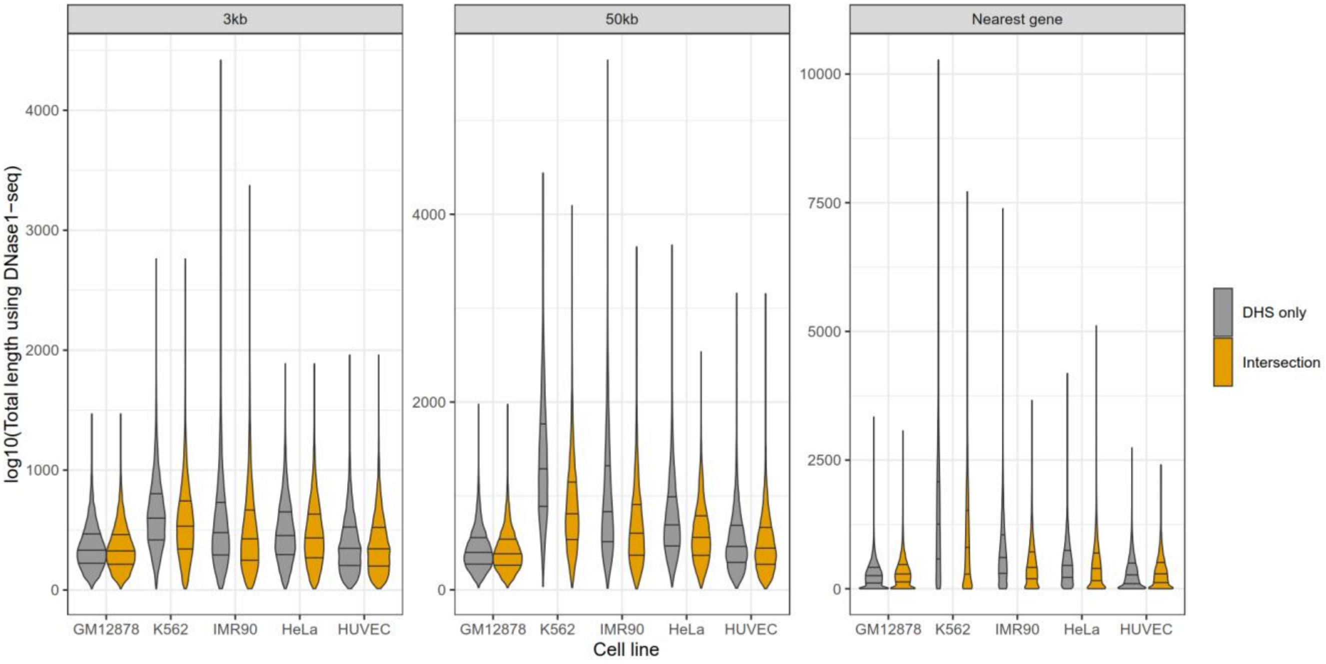


**Supplementary Figure 8:** log10 of the average length of considered segments using DNase1-seq (a) and TF ChIP-seq data (b) using the window based linkage with two different window sizes (3kb,50kb). In grey, the length of purely peak based associations is shown, orange shows the length of the ChromHMM segments and blue the intersection between peaks and ChromHMM promoter/enhancer segments.

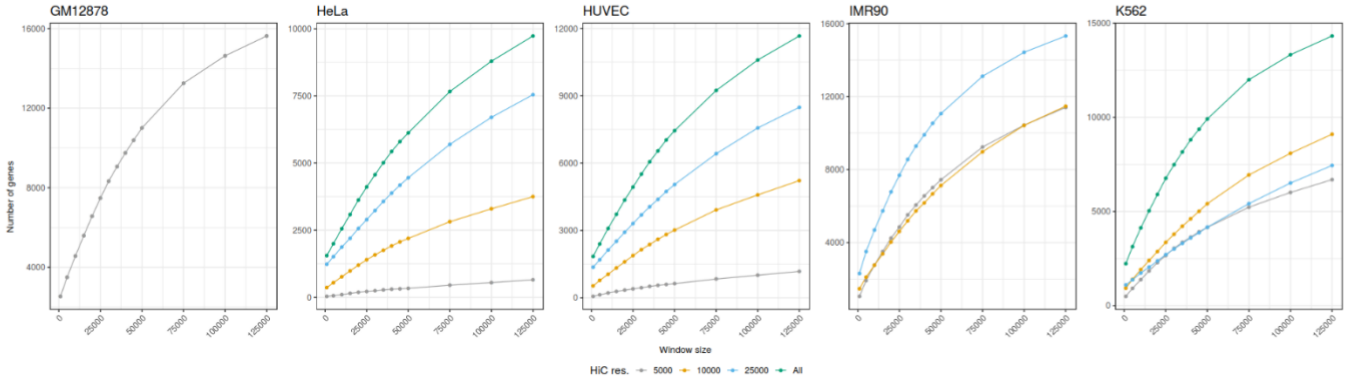


**Supplementary Figure 9:** Here, the relationship between the number of genes overlapping a HiC loop to different HiC resolutions and various loop window sizes is depicted.

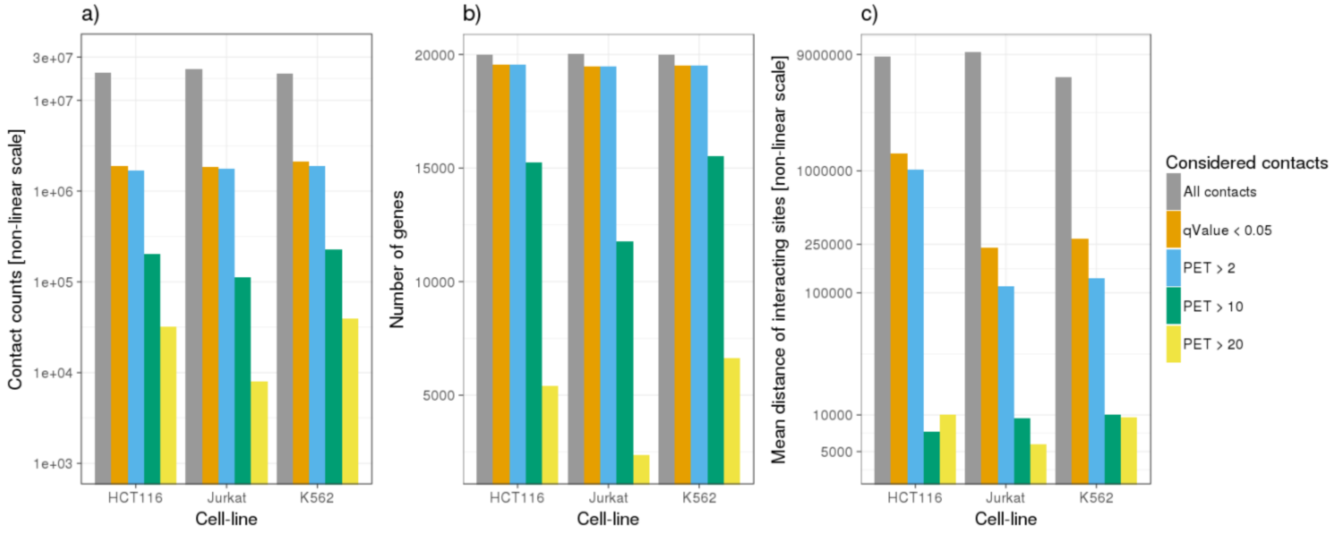


**Supplementary Figure 10:** Filtering of Hi-ChIP data and its influence on (a) contact counts, (b) gene counts and (c) the average distance of the interacting sites.

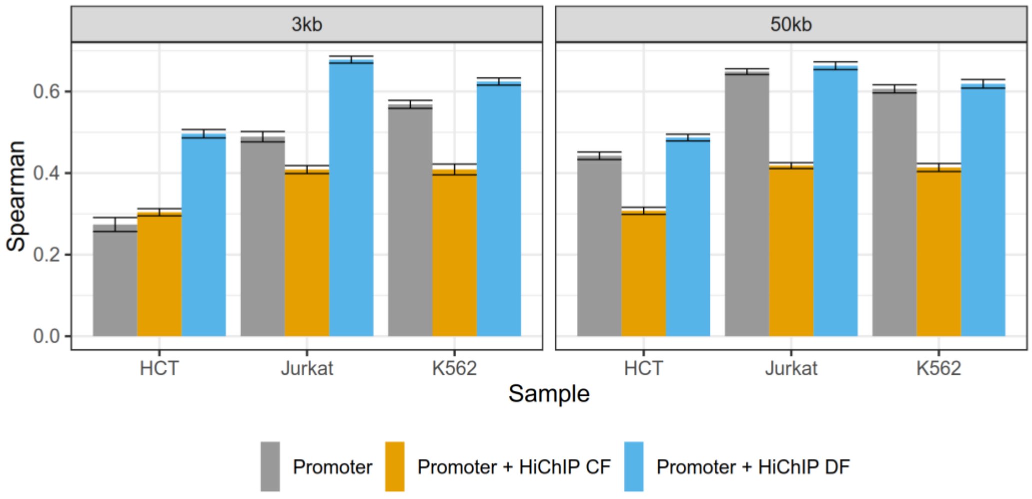


**Supplementary Figure 11:** Here, the performance of spearman correlation obtained for gene-expression prediction models is shown for three different peak feature setups: Promoter only, Promoter and HiChIP peak features combined (CF), Promoter and HiChIP peak features considered separately (DF). Further, we considered two promoter window sizes: 3kb and 50kb.

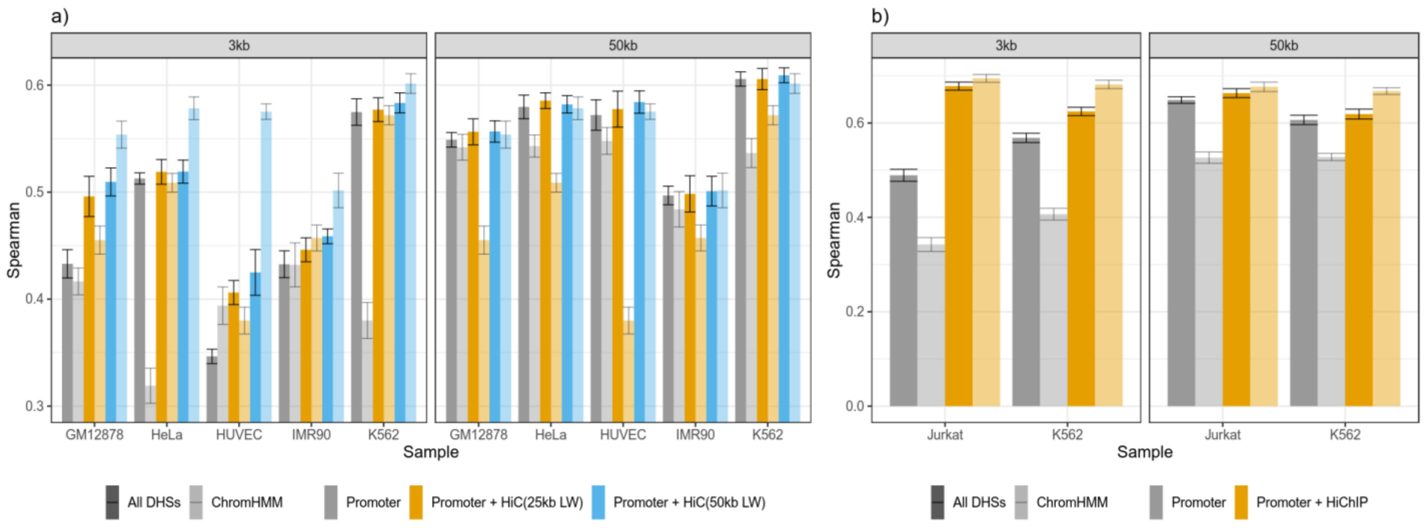


**Supplementary Figure 12:** This figure illustrates the effect of a ChromHMM overlap with a) HiC and b) HiChIP regions on model performance. For HiC and HiChIP models, we consider only the separate feature representation.

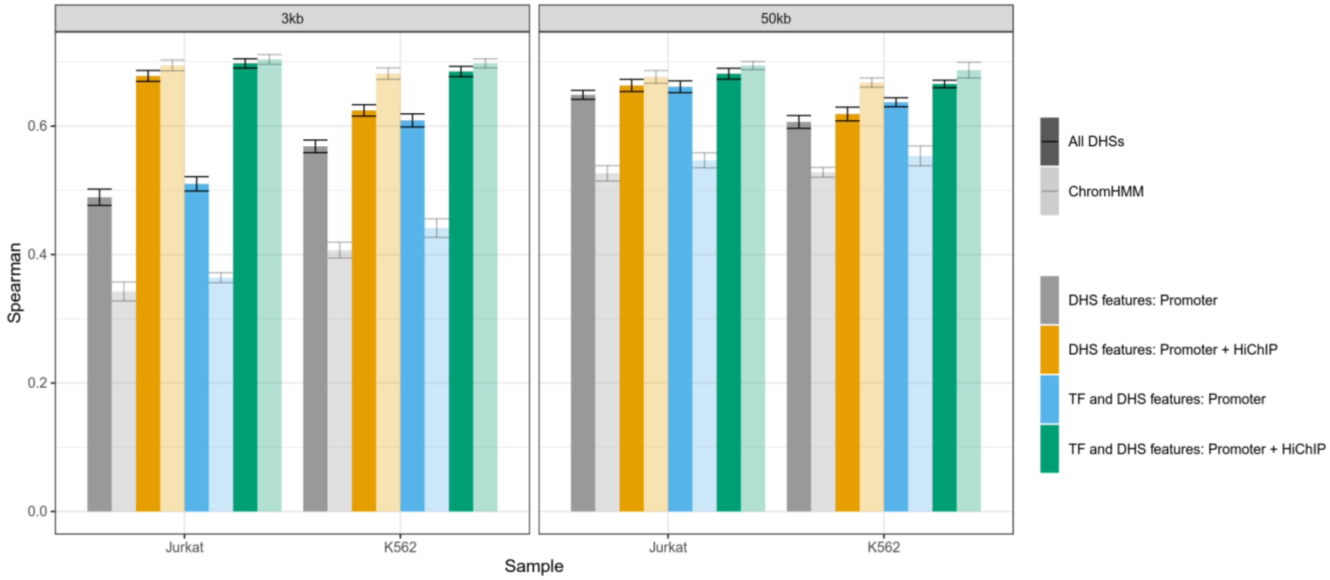


**Supplementary Figure 13:** Performance of gene-expression models in terms of Spearman correlation, using peak and TF affinity features with and without ChromHMM filtering.

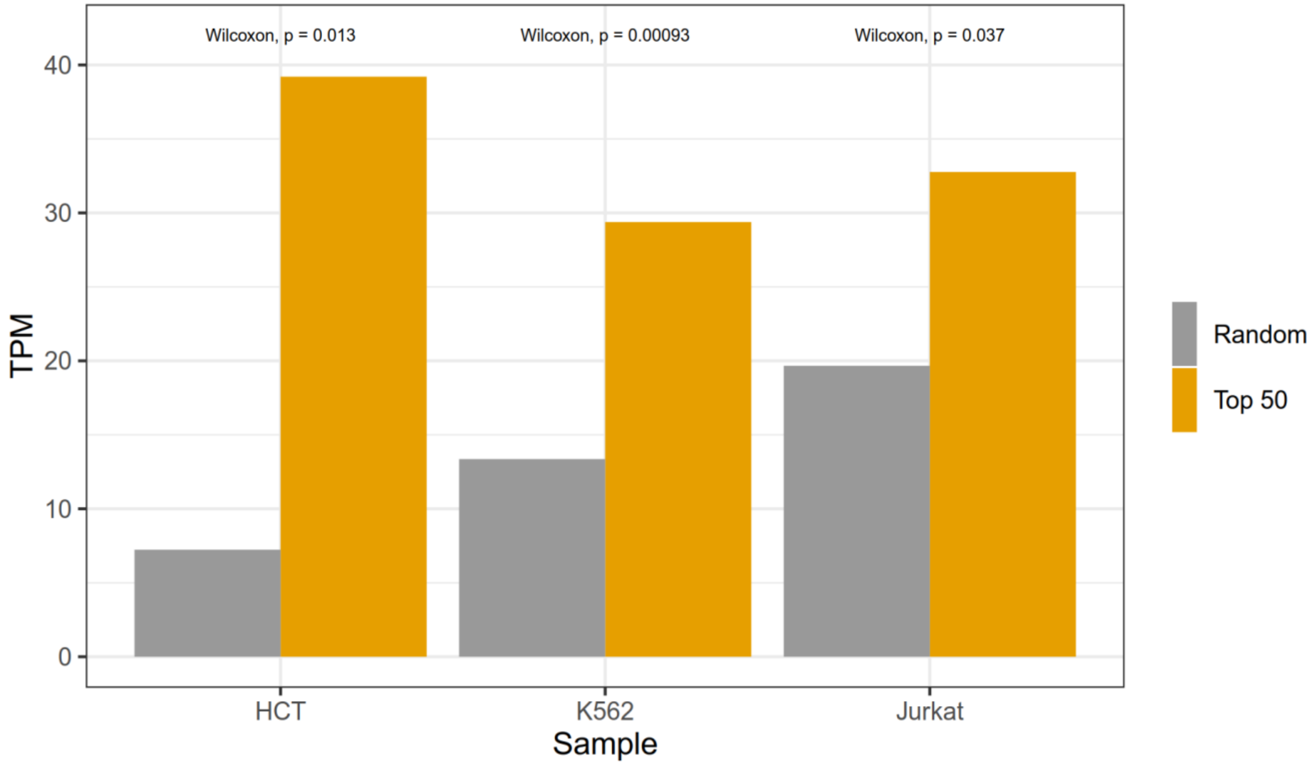


**Supplementary Figure 14:** Expression of TFs measured in TPM for the top 50 TFs and 1000 randomly sampled sets of size 50.

### 3. TEPIC Hi-C extension

We have extended the original TEPIC pipeline with a separate module, to incorporate chromatin conformation capture data, such as HiC and HiChIP data.The new modules incorporates so-called loop list files, which are produced for example by the *HiCCUPS* peak-calling algorithm.

However, we assured that the used format is very simplistic, such that any custom genomic contact information could be used as well. The files are tab separated stating the genomic position of the two loop sites in the following way:

***chr <tab> pos1 <tab>  pos2 <tab> chr <tab> pos1 <tab>  pos2 <tab> <track color> <tab> <contact counts>***

Aside from the loop list file, the Hi-C module offers a parameter controlling the loop window.  It is used to identify loops that interact with the 5' TSS of the gene of interest. Centered at the TSS, the implementation searches for genomic loci marking loop-sites in the loop list.

• **-h** If the name of the Hi-C loop file is provided, all open chromatin regions will be intersected with loop regions around the TSS of each gene.

The optional parameter is:

• **-s** Defines the size of the loop window [bp]. Default is 25000**.**

The double feature space TF affinities are computed automatically if the **-h** parameter is used, unless the **-q** parameter is set as well, which produces peak features only.
